# Molecular mechanisms of recruitment, function and regulation of UPF1 in histone mRNA decay

**DOI:** 10.1101/2025.02.23.639735

**Authors:** Alexandrina Machado de Amorim, Guangpu Xue, Theresa Dittmers, Wenxia He, Sarah Lewandowski, Cecilia Perez-Borrajero, Juliane Bethmann, Nevena Mateva, Clemens Krage, Vidhyadhar Nandana, Janosch Hennig, Henning Urlaub, William F. Marzluff, Sutapa Chakrabarti

**Affiliations:** Institute of Chemistry and Biochemistry, Department of Biology, Chemistry and Pharmacy, Freie Universität Berlin, Takustr. 6, 14195 Berlin, Germany; Integrated Program for Biological and Genome Sciences, University of North Carolina, Chapel Hill, North Carolina 27599, USA; Department of Biochemistry and Biophysics, University of North Carolina, Chapel Hill, North Carolina 27599, USA; Molecular Systems Biology Unit, European Molecular Biology Laboratory (EMBL), 69117 Heidelberg, Germany; Research Group Bioanalytical Mass Spectrometry, Max Planck Institute for Multidisciplinary Sciences, Am Fassberg 11, 37077 Göttingen, Germany; Institute of Chemistry and Biochemistry, Department of Biology, Chemistry and Pharmacy, Freie Universität Berlin, Takustr. 3, 14195 Berlin, Germany; Chair of Biochemistry IV, Biophysical Chemistry, University of Bayreuth, 95447 Bayreuth, Germany

## Abstract

Animal replication-dependent histone mRNAs end in a conserved stem loop (SL) instead of the canonical poly(A) tail present in all other eukaryotic mRNAs. Degradation of the histone SL at the end of the S-phase is initiated by the stem-loop binding protein SLBP and its interplay with the RNA helicase UPF1 and the exoribonuclease 3’hExo. We report direct interactions between SLBP and UPF1 and show that the unstructured SLBP N-terminus wraps around the UPF1 helicase core, contacting it at multiple sites. Although binding of SLBP to UPF1 impedes unwinding activity, it is critical for efficient histone mRNA decay in cells, as unwinding of the SL facilitates degradation by 3’hExo. Here we show that the UPF1-activator, UPF2, binds 3’hExo, and that UPF2-mediated activation of UPF1 overrides the inhibitory effect of SLBP. Our results highlight the intricate network of UPF1-centric protein-protein and protein/RNA interactions that fine-tunes its unwinding activity and orchestrates timely and efficient degradation of histone mRNA.

## Introduction

DNA replication and chromosome segregation are widely considered the two main events leading to cell division. Replication of DNA in eukaryotes must be coordinated with chromatin assembly to ensure maintenance of chromatin structure in the newly formed daughter cells. This entails a rapid upregulation in cellular levels of histone mRNAs in the DNA synthesis phase (S-phase) of the cell cycle, and an equally rapid decrease when cells exit the S-phase to avoid genome instability that arises from accumulation of excess histones [1, 2]. Metazoans achieve cell cycle-coupled expression of specific histone proteins by stringently regulating the levels of the corresponding transcripts [3]. These transcripts are referred to as replication-dependent (RD) histone mRNAs. Metazoan RD histone mRNAs are unusual in that they contain a stem loop (SL) structure at their 3’ end, instead of a poly(A) tail that is present in all other eukaryotic cellular mRNAs [4, 5]. The SL binding protein (SLBP) binds at the 5’ side of the SL immediately after transcription and is essential for formation of the 3’ end [6, 7]. SLBP remains associated with the histone SL RNA until its decay. The RNA-binding domain (RBD) of SLBP does not bear structural similarity to any known RNA binding/recognition motif, and only binds the histone SL, which it recognizes with high affinity and specificity [8, 9]. The SLBP-bound SL is required for every step of histone mRNA metabolism, from 3’ end formation to nuclear export, translation and decay [10].

Decay of RD histone mRNAs at the end of the S-phase requires translation of the mRNA and that the SL is located close to the stop codon. Degradation is initiated at the 3’-end of the histone mRNA by the 3’-5’ exoribonuclease 3’hExo (also known as ERI1) [11, 12]. 3’hExo consists of an N-terminal nucleic acid-binding SAP (<u>S</u>AF-box, <u>A</u>cinus and <u>P</u>IAS) domain and a C-terminal nuclease domain that are connected by a flexible linker. The topology of the nuclease domain classifies it as a DEDDh nuclease that requires two divalent metal ions for catalysis, although the similarity in primary structure to other DEDDh family members is very low [13]. Structural studies of a ternary complex of SLBP:3’hExo:SL RNA showed that the two proteins stably bind the histone SL on opposite sides but do not directly interact with each other [9]. Nevertheless, binding of either factor induces a slight conformational change in the SL, promoting binding of the other. Despite the active nuclease, the SL remains intact in the ternary SLBP-SL-3’hExo complex and is only slightly shortened in the binary SL-3’hExo complex, where the RNA does not reach the nuclease active site. In addition to SLBP and 3’hExo, the RNA helicase UPF1 was also shown to be required for RD histone mRNA decay [14]. An iCLIP analysis of UPF1 occupancy on mRNA showed that UPF1 is recruited to the 3’-untranslated region (UTR) of RD histone mRNA, upstream of the SLBP-binding site [15]. Two lines of evidence suggest a role of UPF1 in RD histone mRNA decay: it was shown to co-immunoprecipitate with SLBP in an RNA-independent manner, and either knockdown of UPF1 or overexpression of a mutant incapable of binding ATP (UPF1 K498A) slows histone mRNA degradation in cells [14].

UPF1, an essential protein in mammalian cells, is involved in many cellular mRNA decay pathways and plays a critical role in the regulation of gene expression [16]. Its role in mRNA decay has been most investigated in the context of nonsense-mediated mRNA decay (NMD), where it was shown to use its RNP-remodelling activity to disassemble mRNPs undergoing NMD [17, 18]. More recently, the ATPase activity of UPF1 was found to be essential for accurate NMD target selection and therefore, critical for progression of NMD [19]. The catalytic activity of UPF1 is stimulated upon binding the core NMD factor UPF2, which induces a large conformational change in the helicase [20, 21]. Binding of UPF2 also promotes UPF1 phosphorylation by the PI3K-like kinase SMG1 and triggers release of RNA from UPF1 [22-24]. As such, UPF2 is not essential for NMD but can modulate it by influencing UPF1-RNA association kinetics and through protein-protein interactions mediated via the exon junction complex (EJC), which recruits the NMD machinery to defective mRNAs [25, 26].

While the function of UPF1 has been extensively studied in NMD, its role in pathways of functional mRNA decay is less understood from a mechanistic point of view. In this study, we present molecular mechanisms for the recruitment, function, and activation of UPF1 in RD histone mRNA decay. We report a direct interaction between SLBP and UPF1 that impedes the unwinding activity of UPF1. Disruption of this interaction leads to a decrease in the rate of histone mRNA decay. Furthermore, we find that the unwinding activity of UPF1 can be stimulated by UPF2 in context of the histone mRNP. Our study shows, for the first time, a direct molecular link between RNA unwinding by UPF1 and RNA degradation in cells and reveals an intricate interplay among key players in the pathway to ensure efficient and timely degradation of RD histone mRNA.

## Results

### UPF1 engages SLBP via direct protein-protein interactions

To analyze the mechanism of recruitment of UPF1 to the histone mRNP, we first investigated the interactions between UPF1 and SLBP. Kaygun and co-workers previously showed that UPF1 co-immunoprecipitates with SLBP within minutes of DNA replication inhibition [14]. The interaction is independent of RNA, suggesting that the association of UPF1 with the histone mRNP is driven by protein-protein interactions. We therefore tested whether UPF1 and SLBP interact *in vitro* using purified proteins. UPF1 is a multi-domain protein, containing the N-terminal cysteine-histidine rich (CH) domain and the helicase core comprising the RecA1 and RecA2 domains and auxiliary domains 1B and 1C (Figure 1A). The structured domains are flanked by long intrinsically disordered regions at the N- and C-termini that harbor serine-glutamine (SQ) motifs which are sites of phosphorylation by the PI3K-like kinase SMG1 [27]. We initially tested three UPF1 proteins which encompass different domains. UPF1-CHh contains both the CH and the helicase core domains, while UPF1-Hel and UPF1-CH contain only the helicase core and the CH domains, respectively (Figure 1A). We performed GST-pulldown assays using GST-tagged UPF1 proteins as baits and full-length SLBP (SLBPfl) expressed and purified from insect cells as the prey. GST-fused to the GYF domain of the translation repression factor GIGYF2 was used as a negative control in this and all other GST-pulldown experiments, unless specified otherwise. GST-UPF1-CHh and UPF1-Hel showed a strong interaction with SLBPfl, suggesting that UPF1 engages SLBP via direct protein-protein interactions mediated by its helicase core (Figure 1B, lanes 1 and 2), independent of SLBP binding to the SL RNA. GST-UPF1-CH showed no binding to SLBP indicating that the CH domain does not participate in this interaction (Figure 1B, lane 3). We next proceeded to map the UPF1-binding region on SLBP by GST-pulldown. SLBP contains a central RBD, flanked by intrinsically disordered regions (IDRs) at the N- and C-termini. We tested four SLBP proteins for their ability to bind UPF1: full-length SLBP (SLBPfl), SLBP-N that contains the N-terminal IDR and the RBD, SLBP-C encompassing the RBD and the C-terminal IDR and the SLBP-RBD alone (Figure 1A). GST-UPF1-Hel was used as a bait in this GST pulldown experiment. We found that GST-UPF1-Hel interacted with SLBPfl and SLBP-N but not with SLBP-RBD or SLBP-C, suggesting that the primary binding site for UPF1 resides in the N-terminal IDR of SLBP (Figure 1C and supplementary figure 1A). SLBPfl and SLBP-N (but not SLBP-C) migrated as doublets on SDS-PAGE gels and were converted to single faster-migrating species upon treatment with calf intestinal phosphatase (CIP) (Supplementary figure 1B), indicating partial phosphorylation of SLBP, likely on Thr 60-61 within the N-terminal IDR [28]. As both the phosphorylated and unphosphorylated SLBP proteins were co-precipitated with UPF1, it appears that phosphorylation does not impact the interaction of SLBP with UPF1. Our observations define the direct protein-protein interactions between UPF1 helicase core and the SLBP N-terminal IDR and are in line with the previously reported RNA-independent interactions between these factors.

**Figure 1.**
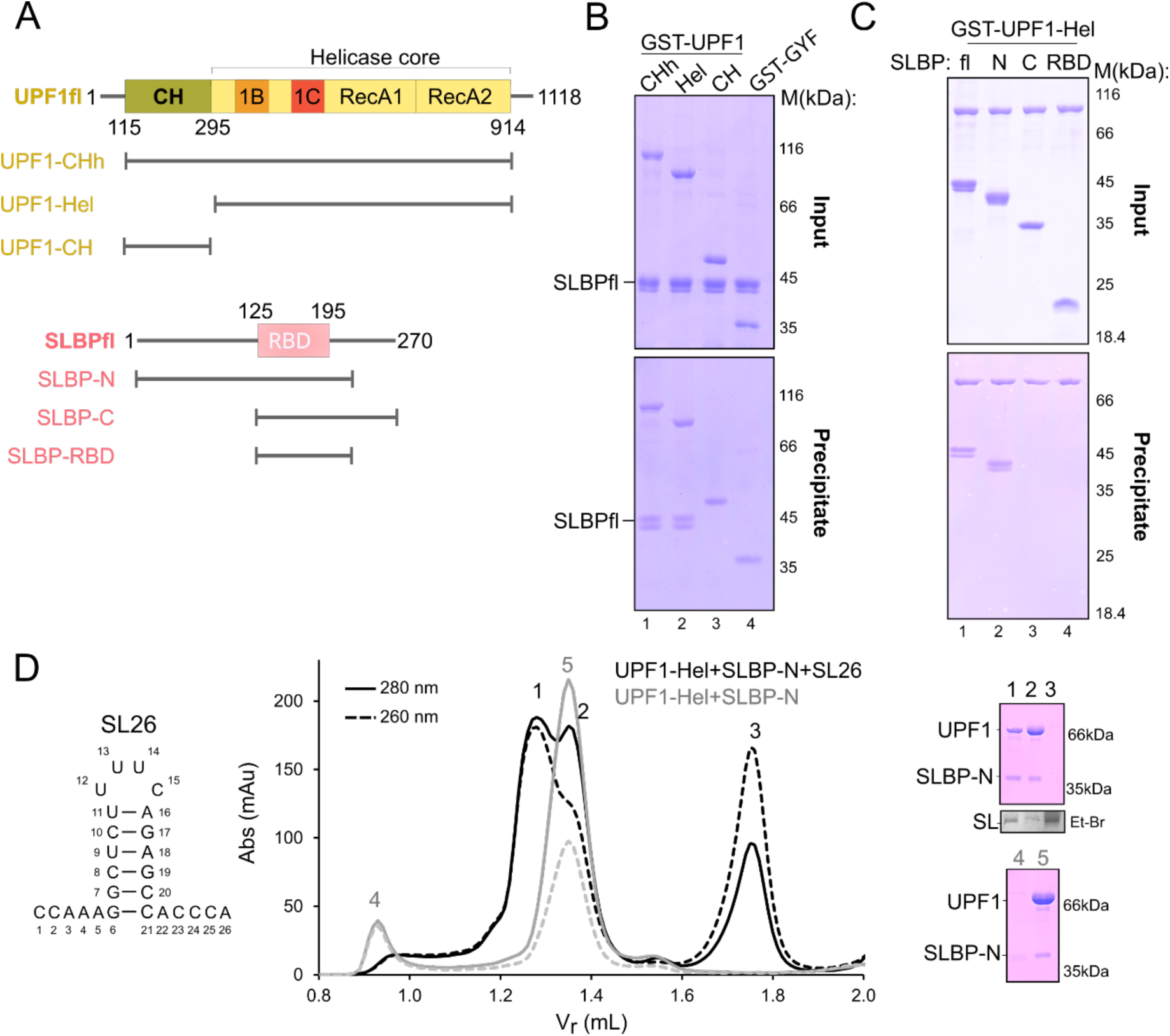
SLBP engages the UPF1 helicase core via direct protein-protein interactions. **(A)** Schematic representation of the domain arrangements of human UPF1 and SLBP. Globular domains are shown as rectangles, flexible linkers and intrinsically disordered regions are shown as lines. Numbers indicate domain boundaries. Protein variants used in this study are shown below each schematic. **(B)** GST-pulldown assays of GST-UPF1 proteins with SLBPfl. GST-GYF serves as negative control. The top and bottom panels correspond to inputs and precipitates in this and all subsequent GST-pulldown experiments, unless specified. The helicase core of UPF1 is sufficient for its interaction with SLBP. **(C)** GST-pulldown using GST-UPF1-Hel as a bait and SLBP variants as preys. A complete gel including negative controls with GST-GYF is shown in Supplementary figure 1A. The N-terminal IDR of SLBP is necessary for binding to UPF1. **(D)** The histone SL RNA stabilizes the UPF1-SLBP interaction. Analytical size exclusion chromatography (SEC) of UPF1-Hel and SLBP-N in the absence (grey traces) and presence (black traces) of the 26 nt-histone SL (SL26). The RNA sequence is shown on the left while PAGE analyses corresponding to the SEC runs are on the right (top: with SL26, bottom: without SL26). The SL-RNA was visualized by ethidium bromide (Et-Br) staining of the urea-PAGE gel.

Like RD histone mRNA, the expression of SLBP protein is also upregulated as cells enter the S-phase of the cell cycle, as part of the mechanism activating histone pre-mRNA processing [29]. SLBP binds the pre-mRNA to recruit U7 snRNP for processing and remains bound to the mature histone mRNA [7, 30]. SLBP is required stoichiometrically since it remains associated with the histone mRNA throughout its lifetime. Since the substrate for histone mRNA degradation is the mRNA-bound to SLBP, we asked whether binding of SLBP to the SL RNA influences its binding to UPF1. To address this, we performed analytical size-exclusion chromatography (SEC) to assess the interaction between UPF1 and SLBP in the absence and presence of the histone SL RNA. The SL RNA used for this analysis (SL26) corresponds to the 26-nucleotide (nt) stem-loop structure generated upon 3’-end processing of the histone pre-mRNA (Figure 1D, left). Analysis of an equimolar mixture of UPF1, SLBP-N and SL26 revealed three peaks, which we analysed by SDS- and urea-PAGE (Figure 1D, middle and right panels). Peak 1 contained stoichiometric amounts of UPF1 and SLBP-N, confirming formation of a stable complex of these components. The ratio of absorbance at 260 nm to that at 280 nm in peak 1 confirms the presence of bound RNA in this complex. Peaks 2 and 3 contained excess UPF1 and SL26 RNA, respectively. The retention volume of peak 1 was lower than that obtained from a mixture of UFP1 and SLBP-N without SL26 (peak 5), consistent with formation of a higher-molecular weight species (Figure 1D, middle panel, compare black and grey traces), containing UPF1 bound to the SLBP-SL26 RNA complex. The retention volume of peak 5 is very similar to that of the individual UPF1 and SLBP proteins, pointing to the absence of a stable complex (Figure 1D and supplementary figure 1C). SL RNA is necessary for SLBP to adopt a correct fold and functional conformation as SLBP-N was partially aggregated in the absence of SL26 (Figure 1D, peak 4). We concluded that although the interaction between SLBP and UPF1 is driven by direct protein-protein interactions, the histone SL plays a key stabilizing role, likely by maintaining SLBP in a functionally competent conformation since the RNA binding domain is not stably folded in the absence of RNA.

### The N-terminal IDR of SLBP engages UPF1 to mediate efficient histone mRNA decay

To gain further insights into the UPF1-SLBP interaction in context of the mRNP, we performed crosslinking mass-spectrometry (CXMS) on an *in vitro* reconstituted ternary complex of UPF1-Hel, SLBP-N and 12U-SL26 RNA (SL26 with an overhang of 12 uridines at its 5’-end), using the long-range crosslinker BS3 (bis(sulphosuccinimidyl)-suberate) that can span lysines 30 Å apart in three-dimensional space (Supplementary Figure 2A). We found multiple crosslinks between the SLBP N-terminal IDR and the RecA domains and domain 1B of UPF1 (Figures 2A and 2B). Although the SLBP-RBD did not bind UPF1 in our biochemical experiments, we observed crosslinks between 2 of the 10 lysines within SLBP-RBD to UPF1 (Supplementary figure 2B). It is possible that the SLBP-RBD bound to the SL-RNA is positioned close to the UPF1 helicase core, without directly interacting with it, and is therefore crosslinked by BS3. Correspondingly, lysine residues of SLBP that directly contact the SL RNA were not crosslinked to UPF1, validating the specificity of the approach in mapping UPF1-SLBP interactions in context of the histone decay mRNP. A similar pattern was observed upon using a lysine-cysteine crosslinker, SMPB (succinimidyl 4-(p-maleimidophenyl)butyrate), although the overall number of crosslinks obtained was lower than with BS3 (Supplementary figure 2C) because of the low abundance of cysteines.

**Figure 2.**
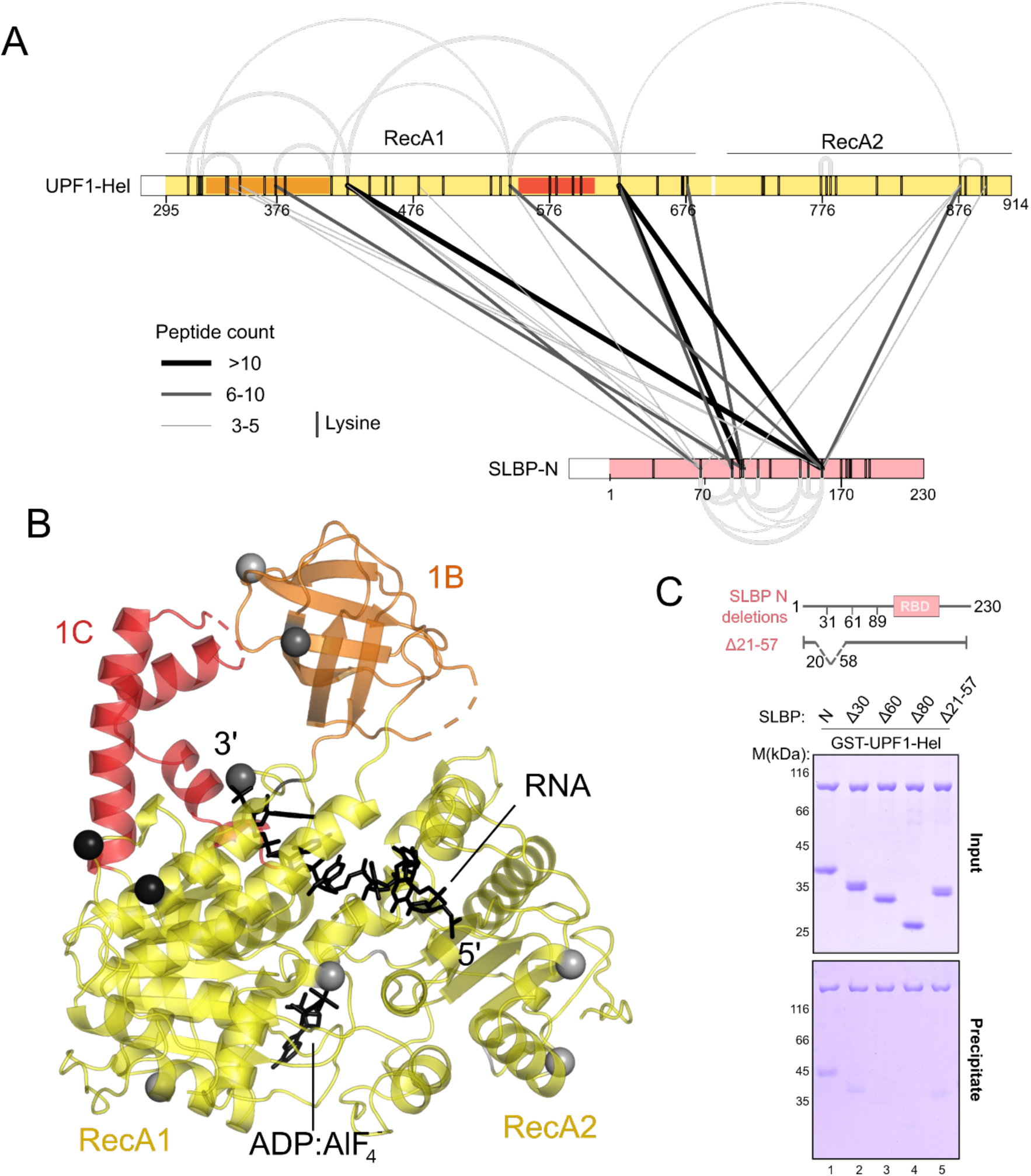
The N-terminal IDR of SLBP binds UPF1 to recruit it upstream of the histone SL. **(A)** Linkage map showing the interlinks between UPF1 and SLBP, derived from crosslinking-mass spectrometry analysis of the UPF1-Hel:SLBP-N:12U-SL26 complex (see Supplementary figure 2A) using BS3. Interlinks are depicted as black or grey lines, where linewidth corresponds to the count of peptides identified for each crosslink. Intralinks within each protein are shown as grey arcs, represented according to peptide count, as for the interlinks. The schematics represent the protein variants used for reconstitution of the complex and are coloured according to Figure 1. Positions of all lysine residues in each protein are indicated by black vertical lines in the schematic. A similar linkage map generated from CXMS analysis using SMPB is shown in Supplementary figure 2C. **(B)** X-ray crystal structure of UPF1-Hel/U_15_ RNA/ADP:AlF_4-_(PDB 2XZO), with the positions of UPF1-SLBP interlinks from (A) highlighted as spheres. The color scheme of the spheres (black, dark grey and light grey) corresponds to the color of the lines denoting interlinks in (A). **(C)** GST-pulldown assay of GST-UPF1-Hel and truncations of SLBP-N. The positions of the N-terminal truncations as well as the deletion within the N-IDR are depicted in the schematic above. Removal of a stretch of 30 residues anywhere within the SLBP N-terminal IDR weaken its binding to UPF1 (see also Supplementary figure 2C). A complete gel including negative controls with GST-GYF is shown in Supplementary figure 2D.

The distribution of UPF1-SLBP interlinks led us to hypothesize SLBP lacks specific interaction motifs, and instead engages UPF1 in a fuzzy interaction, making multiple contacts with the helicase domain through residues located within its N-terminal IDR. To test this hypothesis, we generated a series of SLBP variants, each time truncating approximately 30 residues from the N-terminus (Figure 2C). The truncations were analysed for binding to GST-UPF1-Hel in a GST-PD experiment. We observed a weakening of the SLBP-UPF1 interaction upon deletion of the first 30 amino acids, with a complete loss of binding upon removal of the first 60 amino acids of SLBP. Interestingly, internal deletion of stretches of approximately 30 residues from the N-terminal IDR (amino acids 21-57 or 31-60) also weakened the affinity of SLBP for UPF1 (Figure 2C, Supplementary figures 2D and 2E). Taken together, our results suggest that SLBP uses multiple low-affinity interaction motifs/interfaces to engage UPF1, which likely facilitate its recruitment to the 3’-UTR of histone mRNA, proximal to the histone SL.

### Deletion of exon 2 reduces the rate of histone mRNA degradation

We next proceeded to investigate the effect of binding of UPF1 and SLBP on histone mRNA decay in cells. We took advantage of the fact that the second exon of the SLBP gene (111 nt encoding amino acids 21-59, Figure 3A) could be removed to create an in-frame deletion. A deletion of the second exon in HCT116 cells was engineered using CRISPR/Cas to produce two clonal cell lines (SLBPΔE2, Figure 3A). The SLBPΔE2 cell lines exclusively express the SLBPΔ21-59 protein at levels comparable to full-length SLBP (Figure 3B), and the cells grew normally indicating that this region is not essential. To determine if deletion of this region affects histone mRNA degradation, wild type (wt) and SLBPΔE2 cells were treated with hydroxyurea, an inhibitor of DNA replication, to induce histone mRNA decay and the levels of histone H2a mRNA were monitored by northern blotting (Figure 3C). Wild type cells expressing full-length SLBP showed a strong reduction in H2a transcript levels within 20 minutes of hydroxyurea treatment and >90% depletion of the transcript after 40 minutes. In comparison, H2a mRNA was not rapidly degraded in the SLBPΔE2 cell lines, with as much as 50% of starting amounts remaining after 20 minutes of induction of histone mRNA decay (Figure 3C, right panel). A further depletion of histone mRNA levels was observed after 40 minutes but the amount of H2a mRNA remaining in cells expressing SLBPΔ21-57 was higher than that in wild type cells. We concluded that an interaction between UPF1 and SLBP is crucial for ensuring efficient and rapid degradation of replication-dependent histone mRNA. Nevertheless, weakening the UPF1-SLBP interaction does not entirely block histone mRNA decay, suggesting that this mutation reduced, but did not eliminate, recruitment of UPF1 to the histone mRNA. This is consistent with our previous findings that expression of a catalytically inactive UPF1 protein (UPF1 K498A) slowed down, but did not block degradation of H2a mRNA [14].

**Figure 3.**
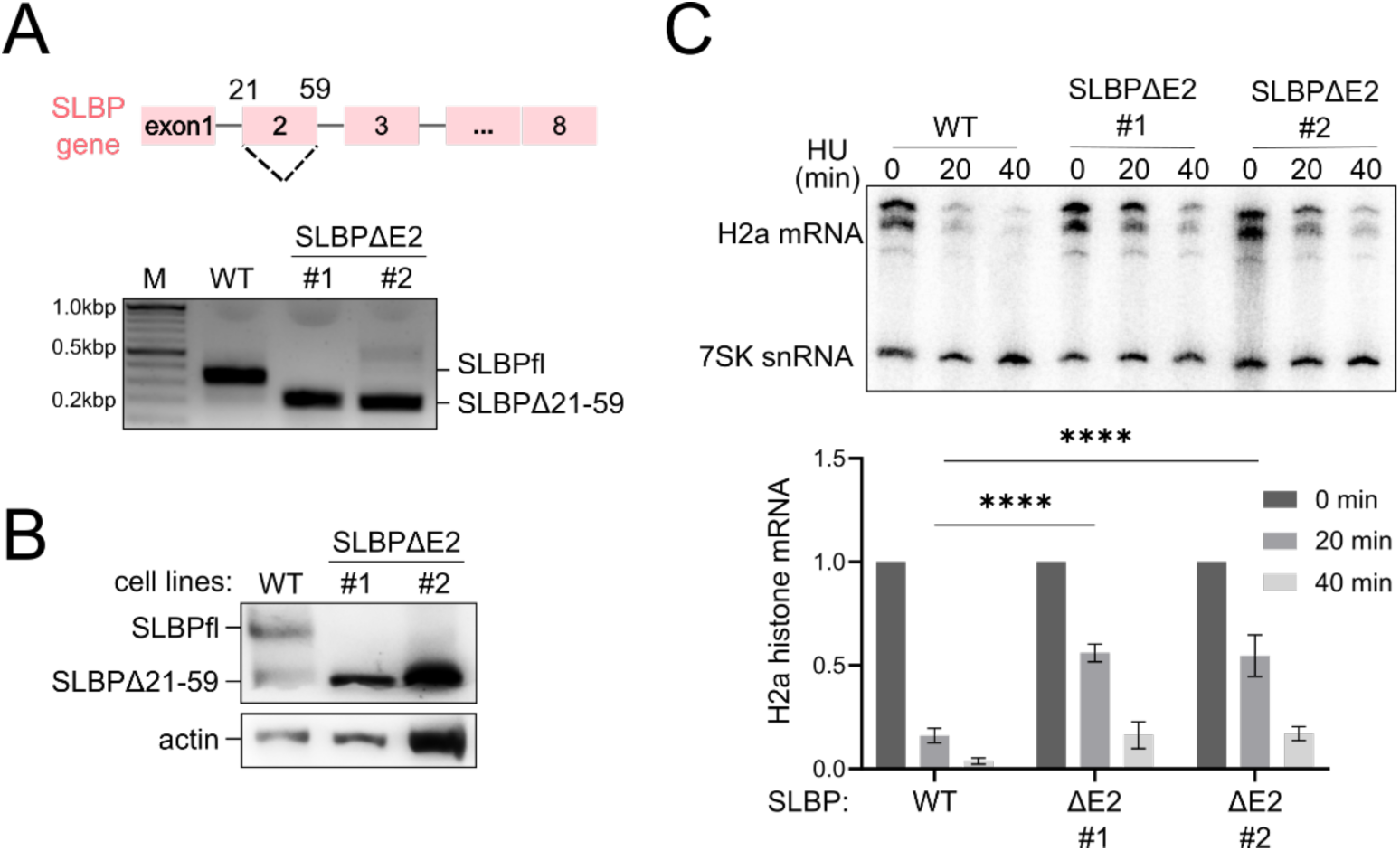
Disruption of the UPF1-SLBP interaction slows down the rate of RD histone mRNA decay in cells. **(A)** Gene organization of the human SLBP gene (top panel). Deletion of exon 2 of the endogenous SLBP gene by CRISPR/Cas corresponds to a deletion of residues 21-59 in the SLBP protein. PCR analysis of the genome locus of the SLBP gene in wildtype and CRISPR/Cas edited HCT116 cells (bottom panel). Primers were designed to amplify a gene fragment flanking exon 2. The amplified fragment in CRISPR/Cas edited cell lines (SLBPΔE2 #1 and #2) are smaller, confirming successful deletion of exon 2 in the endogenous SLBP gene. **(B)** Western blot analysis of wildtype and edited cells lines (SLBPΔE2) using an SLBP-specific antibody shows expression of wildtype and truncated SLBP (Δ21-59). Actin (lower panel) was used as a loading control. A small amount of exon skipping (skipping of exon 2) was observed in wild type cells. **(C)** A weakened UPF1-SLBP interaction significantly reduces the efficiency of histone mRNA decay. Northern blot analysis (top) and corresponding quantification (bottom) of H2a mRNA decay over time, upon hydroxyurea (HU) treatment of wildtype and SLBPΔ2 cell lines. 7SK snRNA served as a loading control. The quantitative data were derived from densitometric analysis of the northern blot and are a representation of the mean values and standard deviation from 3 independent experiments. **** indicates p<0.0001.

### SLBP directly and indirectly impacts unwinding of the histone SL by UPF1

UPF1 is a member of the SF1B family of RNA helicases that translocate on RNA in the 5’-3’ direction to unwind structured RNA or remodel RNPs [31]. Given that the catalytic activity of UPF1 is extensively regulated by intra- and inter-molecular protein-protein interactions, we sought to investigate whether binding of SLBP to UPF1 affects its catalytic activity [20, 32]. To this end, we employed a previously reported fluorescence-based nucleic acid unwinding assay, for which a short fluorescence-labelled DNA strand hybridized to the 3’-end of an unstructured RNA serves as the substrate (linear RNA, Figure 4A) [33]. ATP-dependent displacement of the DNA strand from the RNA:DNA hybrid by the helicase is monitored over time. The constitutively active UPF1 variant, UPF1-Hel, which binds SLBP, was used in these experiments. As previously reported, UPF1-Hel efficiently unwound >75% of the unstructured linear RNA within the first 10 minutes. Addition of an equimolar amount of SLBPfl reduced the extent of nucleic acid substrate unwound by UPF1-Hel to 66% (Figure 4B, compare black and pink traces). Increasing the amount of SLBPfl to 3-fold molar excess of UPF1 led to a further decrease in the fraction of substrate unwound, indicating that binding of the SLBP N-terminal IDR to the UPF1 helicase core impedes its unwinding activity (Figure 4B, brown trace). To ascertain that the observed decrease in unwinding is a consequence of direct SLBP-UPF1 interactions, we measured the unwinding activity of UPF1-Hel in presence of SLBP-RBD which lacks the UPF1-binding region. Addition of up to 3-fold excess of SLBP-RBD did not lead to a significant decrease in the fraction of substrate unwound by UPF1-Hel (Figure 4C). We next sought to analyze the impact of SLBP on UPF1 catalytic activity in context of the histone SL-RNP, and therefore incorporated the histone SL in the RNA strand of the substrate, upstream of the complementary DNA annealing site at the 3’-end (Figure 4A, SL RNA). We hypothesized that UPF1 would translocate along the single-stranded stretch of RNA, encounter and unwind the histone SL, and subsequently the RNA:DNA hybrid at the 3’-end. UPF1-Hel unwinds the histone SL RNA substrate as efficiently as the linear RNA substrate, showing that the 6 bp stem is not sufficient to impede the unwinding activity (Supplementary figure 3A). As observed with the linear RNA substrate, addition of SLBPfl, but not SLBP-RBD, leads to an overall decrease in the fraction of SL RNA substrate unwound by UPF1-Hel (Figure 4D). Interestingly, addition of SLBPfl as well as SLBP-RBD appears to decrease the initial rate of unwinding of the SL-substrate by UPF1-Hel (Figure 4D, inset). This effect is not observed with the linear RNA substrate (Figure 4E), suggesting that binding of SLBP to the histone SL modulates the rate of unwinding of the SL RNA substrate by UPF1. We postulate that when bound to the SL-RNA, SLBP creates a roadblock for translocation, and must be actively displaced by UPF1 to reach the 3’-end and unwind the RNA:DNA hybrid. Accordingly, addition of SLBP-RBD only slows down the initial unwinding of the SL RNA substrate but does not affect the fraction of substrate unwound by UPF1-Hel at later time-points. Taken together, our data suggest that SLBP uses two distinct mechanisms to modulate unwinding of the histone SL by UPF1: directly, by binding the UPF1 helicase core, and indirectly, via the strong association of the RBD with the histone SL.

**Figure 4.**
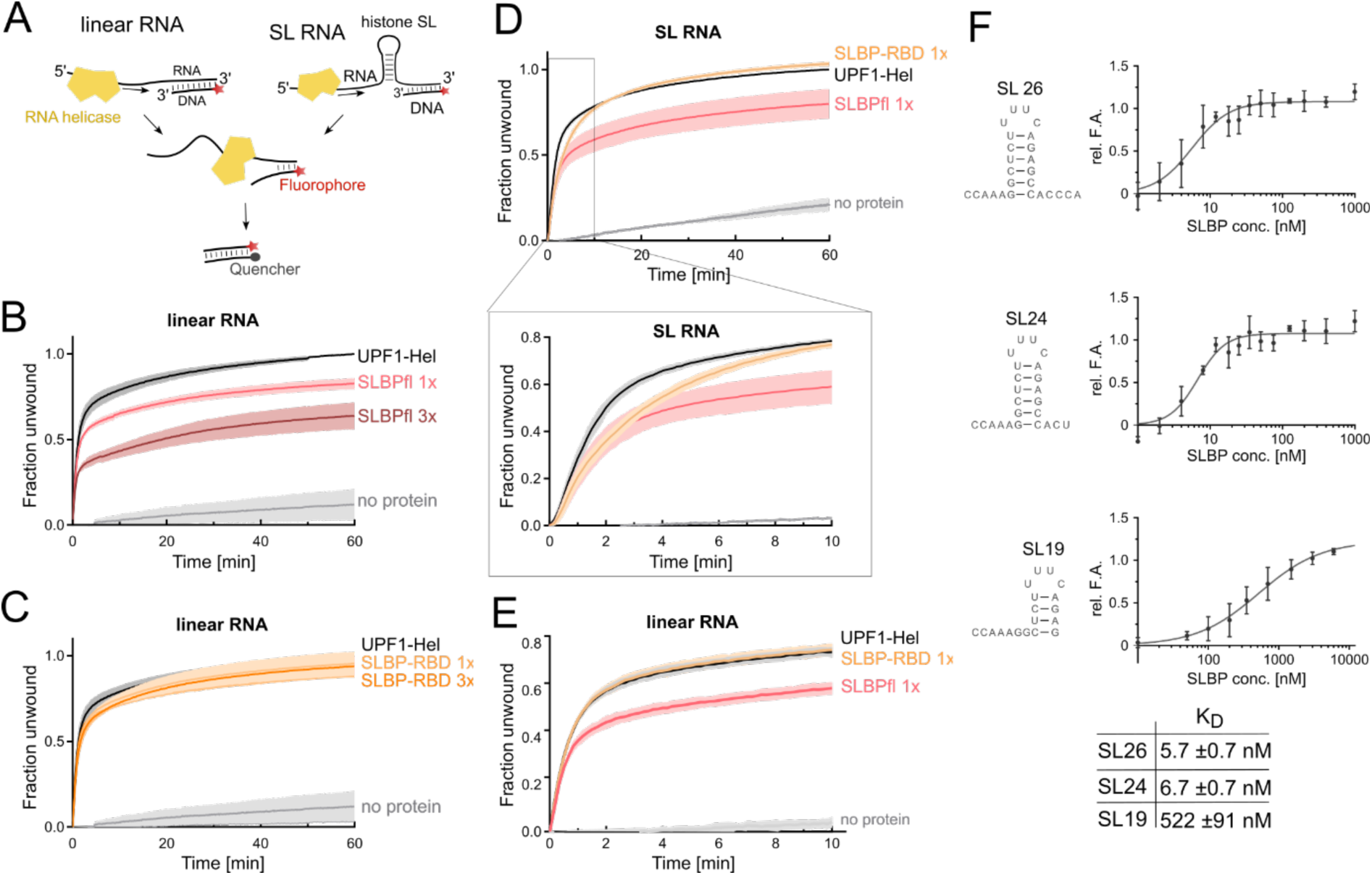
SLBP directly and indirectly impacts unwinding of the histone SL by UPF1. **(A)** Schematic of the experimental setup of the nucleic acid unwinding assay. The RNA:DNA template labelled with Alexafluor 488 on the short DNA strand is the substrate for unwinding. SL RNA and linear RNA refer to substrates containing or lacking the histone SL in the RNA strand of the hybrid, respectively. The DNA strand released upon unwinding of the substrate by UPF1 is trapped by a complementary DNA labelled with a quencher (BHQ1). **(B-C)** Unwinding activity of UPF1hel on the linear RNA substrate in presence of SLBPfl (B) or SLBP-RBD (C). The SLBP proteins were added in equimolar amounts (1x) or in 3-fold molar excess (3x). The fraction of substrate unwound was plotted against reaction time. The individual data points, collected at 10s intervals, represent the mean of at least 3 independent experiments, with technical duplicates for every experiment. Shaded areas represent the standard deviation of each measurement. SLBPfl has an inhibitory effect on the unwinding activity of UPF1. **(D)** Unwinding activity of UPF1-Hel on the SL RNA substrate in the presence of equimolar amounts of SLBPfl (pink trace) and SLBP-RBD (orange trace). The inset below shows an enlarged view of the first 10 minutes of the reaction. SLBP-RBD retards unwinding of the SL RNA substrate by UPF1 in the early stages of the reaction. **(E)** Unwinding activity of UPF1-Hel on the linear RNA substrate in the presence of equimolar amounts of SLBPfl and SLBP-RBD (pink and orange traces, respectively). Only the first 10 minutes of a 60-minute reaction are shown for clarity. SLBP-RBD has no effect on unwinding of the linear RNA substrate. **(F)** Quantitative measurement of the binding affinities of SLBPfl for the histone SL and distinct SL degradation intermediates. The sequence and secondary structure of each RNA is shown on the left. Data points and error bars in each plot represent the mean and standard deviation of at least 3 independent experiments. The dissociation constant (*K*_D_) was determined by fitting the data to an equation representing one-site specific binding with Hill slope (fit denoted by a solid line). The *K*_D_ (and standard deviation) of SLBPfl for each SL RNA is reported in the table below. The binding affinity of SLBP for a partially truncated histone SL (SL19) is 100-fold lower than that for an SL with an intact stem (SL26 and SL24). A qualitative assessment of binding of SLBPfl to SL26, SL24 and SL19 by EMSA is shown in Supplementary figure 3B.

Previous studies on SLBP and the histone SL showed that SLBP binds the 26 nt-SL RNA with very high affinity, rationalizing its role as a roadblock to translocation of UPF1. The mature SL is trimmed by 2-3 nts in the cytoplasm by 3’hExo and oligouridylated by the terminal uridyl transferase, TUT7 [34-37]. The two opposing reactions maintain the SL at a length of ∼24 nt in the cytoplasm over the S-phase of the cell cycle [9, 12]. With the onset of histone mRNA decay at the end of the S-phase, 3’hExo degrades the SL RNA in a distributive manner, leading to accumulation of distinct SL RNA intermediates [38, 39]. To glean further insights into how SLBP acts as a roadblock for UPF1 unwinding and its impact on SL RNA degradation, we set out to determine the binding affinity of SLBP to SL RNA variants using fluorescence anisotropy. The intact SL RNA (SL26), the cytoplasmic variant (SL24) and a stable degradation intermediate lacking 7 nt from the 3’-end (SL19) labelled with 6-FAM at the 5’-end were analysed for binding to SLBP (Figure 4F). The SL19 intermediate was identified by End-Seq analysis of S-phase cells [36, 39, 40]. Consistent with previous reports, SLBP binds SL26 with a very high affinity (*K*_D_ of ∼6 nM) [8]. The cytoplasmic SL, SL24, which contains an intact stem also shows very strong binding (*K*_D_ of ∼7 nM) to SLBP. In contrast, the affinity of SLBP for the intermediate SL19 is drastically reduced (*K*_D_ of ∼500 nM), suggesting that once the SL is partially degraded, SLBP can no longer rebind the intermediate (see also Supplementary figure 3B). We argue that this is likely an important event as SLBP also acts as a roadblock for degradation by 3’hExo in our *in vitro* degradation analysis (Supplementary figure 3C).

### Unwinding of the histone SL by UPF1 enhances the efficiency of histone mRNA decay

To investigate the role of UPF1 in mediating histone mRNA decay at a mechanistic level, we adopted a biochemical approach where we compared the 3’-5’ degradation of a histone SL RNA by the exoribonuclease 3’hExo in the absence and presence of UPF1. The RNA substrate (60N-SL19) used in this setup contains a shortened stem (corresponding to SL19) and a 60-nt 5’-overhang corresponding to the 3’-UTR stretch preceding the SL in H2bc (Supplementary figure 4A). To prevent heterogeneity of the 3’-end length/sequence, we produced this substrate as an HDV ribozyme-fusion by *in vitro* transcription. Autocleavage of the precursor RNA by HDV ribozyme releases 60N-SL19, ensuring identical 3’-ends in all substrate molecules (Supplementary figure 4B). Although 3’hExo binds the SL19 RNA with an affinity comparable to that for SL26 and SL24, our initial studies showed that 60N-SL19 is more amenable to degradation by 3’hExo than 60N-SL24 (Figure 5A and supplementary figure 4C). We reconstituted the substrate with 3’hExo (and UPF1-Hel, where indicated) and added ATP/MgCl_2_ to initiate the reaction. In absence of UPF1, we observed degradation starting from 10 minutes after initiation of the reaction, and a steady progress of the reaction up to 60 minutes. Although the initial substrate was significantly depleted by 60 minutes, it was not completely degraded, evident from the accumulation of stable intermediates (Figure 5B, left panel, lanes 7-12). Addition of UPF1-Hel increased the rate of the reaction but did not change the intermediates generated. Accumulation of the first intermediate was observed as early as 10 minutes and progressed beyond the intermediate obtained in absence of UPF1 at 20 minutes (Figure 5B, left panel, lanes 13-18). UPF1 enhanced the efficiency of degradation by 3’hExo by as much as 30-40% in the later time points of the reaction (Figure 5B, right panel). Interestingly, an identical pattern of degradation was observed upon addition of higher amounts of 3’hExo (Figure 5B, lanes 1-7), supporting the idea that UPF1 facilitates degradation of the histone SL but is not necessary for enabling it. Addition of SLBP did not interfere with the degradation of 60N-SL19 by 3’hExo alone but negated the stimulatory effect of UPF1 on degradation (Supplementary figure 4D). This is consistent with the observations that SLBP does not bind SL19 and therefore does not act as a roadblock for degradation, but that SLBP has an inhibitory effect on UPF1 unwinding activity. Our results suggest that the unwinding activity of UPF1 enhances the efficiency of degradation of histone mRNA.

**Figure 5.**
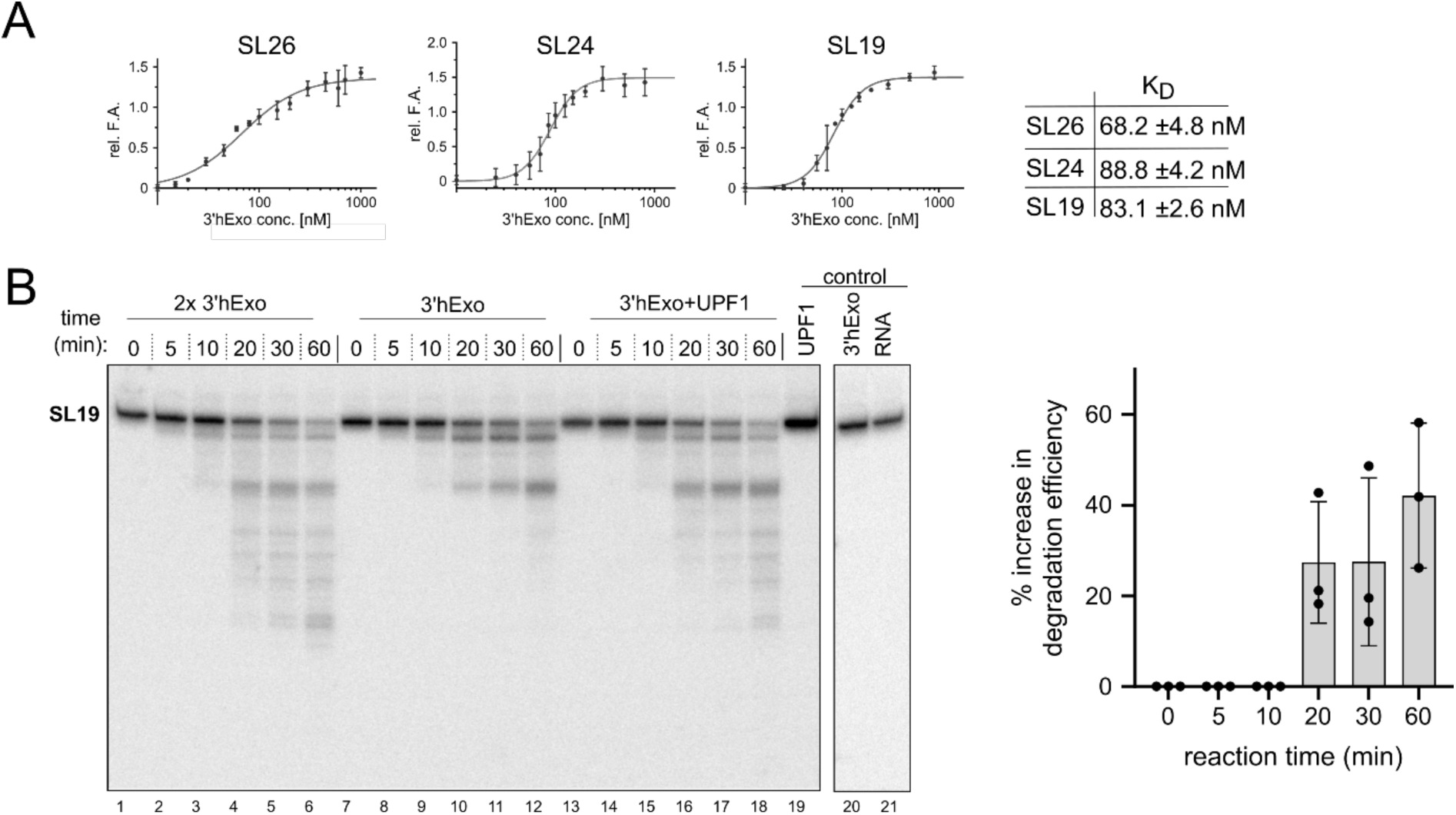
Unwinding of the histone SL by UPF1 enhances the efficiency of histone mRNA decay. **(A)** Fluorescence anisotropy measurements of binding of 3’hExo to SL RNA and distinct SL degradation intermediates to derive dissociation constants (*K*_D_). The data were plotted and *K*_D_s derived as described for Figure 4F. Unlike SLBP, 3’hExo binds all three SL RNA substrates with comparable affinities. **(B)** Time-dependent analysis of degradation of 60N-SL19 RNA by 3’hExo in the absence (lanes 7-12) and presence (lanes 13-18) of UPF1-Hel. As a comparison, 2-fold excess of 3’hExo was used to mimic a more efficient degradation reaction (lanes 1-6). Reactions were initiated by addition of ATP and magnesium ions. Samples were removed from the reaction at the indicated time points, quenched with RNA loading dye and analysed by urea-PAGE and phosphorimaging. Degradation of SL19 by 3’hExo leads to the accumulation of distinct degradation intermediates. Addition of UPF1 increases the efficiency (rate and progression) of the degradation reaction, generating a pattern that is comparable to addition of 2x-3’hExo. Densitometric analysis on the urea-PAGE gel was performed to determine the percentage increase in degradation efficiency upon addition of UPF1 (right panel). The data points and error bars of the plot represent mean values and standard deviation derived from analysis of 3 independent experiments. An increase in degradation efficiency in the presence of UPF1 is observed ∼20 minutes after initiation of the reaction.

### An acidic linker in UPF2 stably associates with 3’hExo

The role of UPF1 unwinding activity and the impact of its regulation by SLBP on histone mRNA decay led us to ask if the core NMD factor UPF2 also plays a role in regulating UPF1 catalytic activity in context of the histone mRNA. The domain structure of UPF2 is shown in Figure 6A. It contains three *m*iddle-of-e*IF4G* (MIF4G) domains and a C-terminal UPF1-binding domain (U1BD) [21, 41]. Binding of UPF2 to UPF1 recruits it to the exon junction complex and stimulates UPF1 catalytic activity by inducing a conformational change in the helicase [20, 21, 24]. Recent studies show that binding of UPF2 to UPF1 drastically reduces the affinity of the helicase for RNA and that UPF2 cannot remain stably associated with RNA-bound UPF1 [23]. It therefore follows that if UPF2 activates UPF1 on the histone mRNA, it must interact with another protein factor within the mRNP, allowing it to be in proximity to the helicase and engage it when necessary. To investigate this possibility, we performed GST-pulldowns of GST-UPF2s with SLBP and 3’hExo. The truncated UPF2s protein spans the domains MIF4G3 and U1BD, which mediate all known interactions of UPF2 (Figure 6A) [21, 42-44]. UPF2s shows a strong interaction with 3’hExo but not SLBP (Figure 6B, lanes 1-2), suggesting that it might be recruited to the histone mRNP via 3’hExo. To map the 3’hExo-binding site on UPF2, we tested interactions of a series of UPF2 constructs with GST-3’hExo (Figure 6C). As expected, we saw a strong interaction of GST-3’hExo with UPF2L and UPF2s, but not with UPF2-MIF4G1-2, supporting our earlier observation that the binding side for 3’hExo resides in the C-terminal region of UPF2 (Figure 6C, lanes 1-3). Systematic truncations of the C-terminal region of UPF2 revealed that the U1BD is not involved in mediating interactions with 3’hExo (Figure 6C, lane 6). The MIF4G3 domain of UPF2 is connected to its U1BD via a stretch of predominantly acidic residues that we refer to as the acidic linker (AL). UPF2 fragments harboring the AL but lacking either the U1BD or the MIF4G3 domains (UPF2 MIF4G3-AL and UPF2 AL-U1BD, respectively) showed strong binding to 3’hExo, suggesting that the 3’hExo-binding site is located within the AL of UPF2 (Figure 6C, lanes 3, 4 and 5). To corroborate these observations, we reconstituted a complex of 3’hExo with the UPF2-AL by size-exclusion chromatography (Figure 6D). While the AL engaged 3’hExo in a stable complex, a UPF2 variant lacking the AL (UPF2ΔAL) failed to interact with 3’hExo (Supplementary figure 5A). We concluded that the AL of UPF2 is necessary and sufficient for mediating a stable interaction with 3’hExo. Reconstitution of a stable ternary complex of 3’hExo-UPF2-UPF1 shows that UPF2 can simultaneously engage with 3’hExo and UPF1 in solution, and that binding of 3’hExo to the UPF2-AL does not perturb the interactions between UPF2-U1BD and UPF1 (Figure 6E).

**Figure 6.**
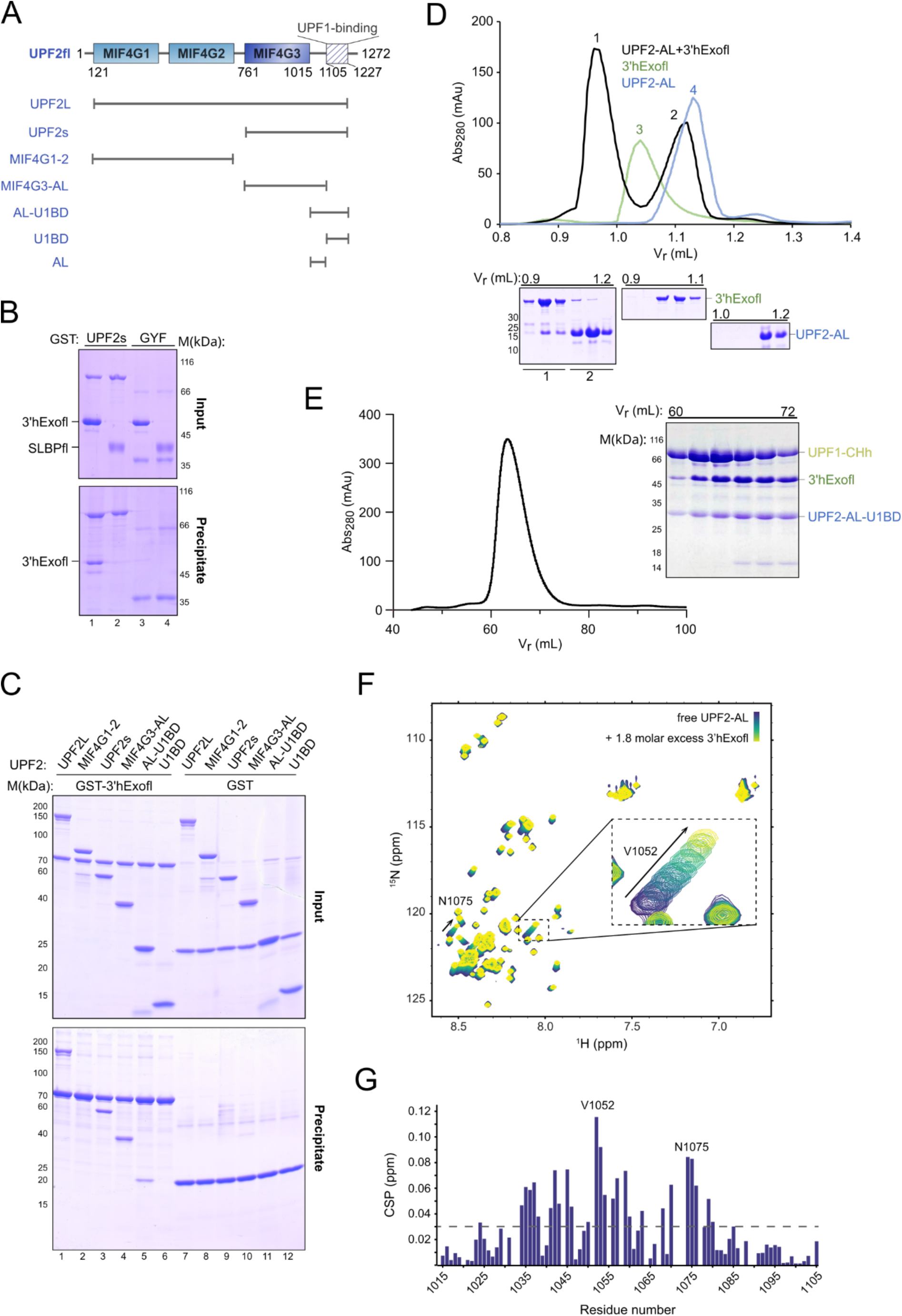
UPF2 stably associates with 3’hExo via an acidic linker at its C-terminal end. **(A)** Schematic representation of the domain organization of human UPF2. The structured middle-of-eIF4G (MIF4G) domains are shown as filled rectangles while the C-terminal U1-binding domain (U1BD) is depicted by a crosshatch. Numbers indicate domain boundaries. Protein variants used in this study are shown as black lines. **(B)** GST-pulldown assay of GST-UPF2s (and GST-GYF as negative control) with full-length SLBP and 3’hExo. 3’hExofl shows a specific interaction with UPF2s while SLBPfl shows no appreciable binding. **(C)** GST-pulldown assays using GST-3’hExofl as a bait and different UPF2 variants as preys show that all UPF2 proteins encompassing the acidic linker (AL) are co-precipitated by 3’hExo. **(D)** Analytical SEC of mixture of UPF2-AL and 3’hExo (black trace) confirms that the UPF2-AL is sufficient for formation of a stable complex with 3’hExo. SEC runs of the individual proteins 3’hExofl (green trace) and UPF2-AL (blue trace) are shown for comparison. Appearance of a peak at lower retention volume relative to the individual proteins in the UPF2-AL:3’hExo mixture indicates stable complex formation. The corresponding SDS-PAGE analysis are shown on the right. See also Supplementary figure 4D. **(E)** Size-exclusion chromatography and corresponding SDS-PAGE analysis depicting formation of a stable ternary complex of UPF1-CHh, UPF2s and 3’hExo. The exclusion volume of the column is 40 mL. **(F)** ^1^H-^15^N-HSQC NMR titration experiments of ^15^N-labeled UPF2-AL with increasing concentrations of 3’hExofl. The spectrum of free UPF2-AL is in blue while those recorded in presence of increasing concentrations of 3’hExofl are in progressively lighter shades of green. Two significant peak shifts are indicated by arrows. The inset shows a zoomed-in view of residue V1052 of UPF2-AL, exhibiting the largest CSP upon addition of 3’hExo. **(G)** Histogram of the chemical shift perturbations (CSP) of UPF2 AL upon titration of 3’hExofl plotted against the UPF2-AL sequence. The dashed line corresponds to the average CSP value obtained.

We next performed GST-pulldowns of 3’hExo truncations, shown in Supplementary figure 5B, with the GST-UPF2s protein. The SAP and nuclease domains co-precipitated with UPF2, suggesting that both domains interact with UPF2-AL (Supplementary figure 5C). To gain insights into the UPF2-3’hExo interaction at a molecular level, we used NMR spectroscopy to monitor the changes in the amide peaks of ^15^N-labelled UPF2-AL upon addition of unlabeled 3’hExofl. We observed chemical shift perturbations (CSPs) in the fast exchange regime in the NMR timescale, indicative of relatively weak binding (Figure 6F). To locate the regions of UPF2-AL most affected in the presence of 3’hExo, we assigned its backbone resonances (^13^C^α^, ^13^C^β^, ^1^H^N^, and ^15^N) and plotted the amide CSPs as a function of residue number (Figure 6G). The largest CSPs were found in a broad region encompassing residues 1030 to 1090 enriched in glutamate residues, indicating that binding did not occur through a highly localized interface of UPF2. The chemical shifts observed also confirmed that UPF2-AL is intrinsically disordered in solution (Supplementary figure 5D) and does not become ordered upon addition of 3’hExo, as shown by the lack of amide dispersion in the bound state. Upon titration of the individual SAP and nuclease subdomains of 3’hExo into ^15^N-labeled UPF2-AL, we observed similar regions being perturbed (Supplementary figure 5E). Therefore, UPF2 interacts with distinct regions of 3’hExo using a similar interface, without making domain-specific interactions. We speculate that a basic groove spanning the SAP and nuclease domains of 3’hExo serves as a docking platform for UPF2-AL, engaging it in low affinity but high avidity interactions that allows formation of a stable complex in solution. Using surface plasmon resonance, we determined the dissociation constant (*K*_D_) of the 3’hExo-UPF2 interaction to be ∼170 μM, in agreement with the *K*_D_ of ∼200 μM estimated from our NMR titrations (Supplementary figures 5F and 5G).

### Activation of UPF1 in context of the histone mRNP

As a first step towards understanding the effect of UPF2 on UPF1 activation in context of the histone SL-RNP, we tested the unwinding activity of UPF1-CHh on the SL RNA substrate. As expected, the rate of unwinding of SL-RNA by UPF1-CHh is lower than that by UPF1-Hel, with only ∼40% of the substrate unwound in the first 10 minutes by UPF1-CHh in contrast to ∼75% by UPF1-Hel (Figure 7A, compare black and grey traces). Consistent with previous studies, addition of UPF2s significantly enhances the unwinding activity of UPF1 (Figure 7A, compare black and blue traces). To dissect the effects of SLBP on UPF2-activated UPF1, we measured the rate of unwinding of the SL RNA substrate by UPF1-CHh in presence of SLBPfl, with and without addition of UPF2s. As described earlier, SLBP has an inhibitory effect on UPF1 unwinding activity, which we attribute to the interaction of the SLBP N-terminal IDR with the helicase core. Addition of SLBPfl also retards the rate of unwinding by UPF1-CHh, which is overridden in the presence of UPF2 (Figure 7B, compare black, red and purple traces). We therefore concluded that UPF2 can activate the UPF1 helicase on the histone SL-RNA.

**Figure 7.**
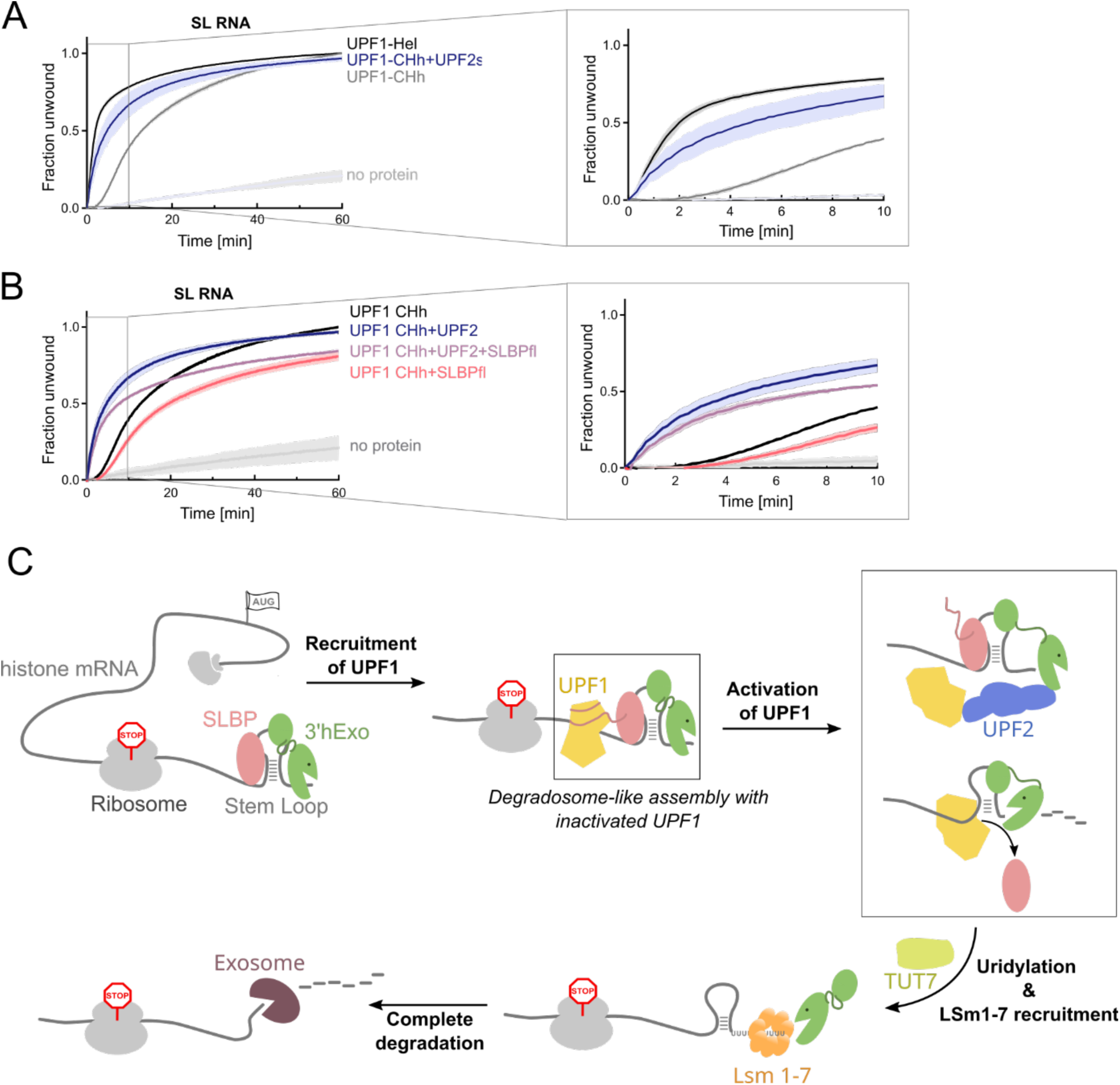
Activation of UPF1 in context of the histone mRNP. **(A)** Comparison of the unwinding activity of UPF1-CHh on SL RNA substrate in the absence (grey trace) and presence of UPF2s (blue trace). The unwinding activity of UPF1-Hel is shown as a reference (black trace). The inset shows an enlarged view of the first 10 minutes of the reaction, where the differences in unwinding activity are the largest. **(B)** Unwinding activity of UPF1-CHh on the SL RNA substrate in the presence of either UPF2 (blue trace) or SLBPfl (red trace) or both UPF2 and SLBP (purple trace). The activity of UPF1-CHh in absence of UPF2 and SLBPfl (black trace) has been shown for comparison. The inset shows an enlarged view of the first 10 minutes of a 60-minute reaction. Activation of UPF1 by UPF2 supersedes the inhibitory effect of SLBP on the helicase. **(C)** Model depicting the recruitment, function and regulation of UPF1 in the context of histone mRNA decay. UPF1 is recruited to the histone mRNA to formation of a degradosome-like assembly with SLBP and 3’hExo in the early stage of decay. Activation of UPF1 by UPF2 leads to displacement of SLBP, unwinding of the SL and degradation by 3’hExo. Initial decay by 3’hExo primes the SL RNA for oligouridylation, mediated by TUT7 and rapid mRNA decay, brought about by binding of the LSm1-7 complex and recruitment of bulk mRNA decay factors.

## Discussion

Like NMD, translation is also required for histone mRNA degradation, and translation termination of histone mRNA has been postulated to be affected by inhibition of DNA replication [45]. Furthermore, UPF1 is also recruited to sites of inefficient translation termination in NMD, implying a similar mode of recruitment to the histone mRNP [46]. It appears that a bipartite signal is required to recruit and activate UPF1 for histone mRNA decay: the paused ribosome on the termination codon and binding to SLBP. Moving the histone SL too far from the termination codon prevents regulated histone mRNA degradation, just as deletion of the UPF1-binding site on SLBP slows down degradation [45].

The studies presented here provide the first mechanistic insights into the molecular network necessary for initiation of regulated histone mRNA decay. Previous reports describe a requirement for the RNA helicase UPF1 in RD-histone mRNA decay [14]. It was shown that 3’hExo, when bound to an SL RNA-SLBP complex cannot degrade beyond two nucleotides from the 3’ end [9]. Identification of distinct decay intermediates resulting from 3’hExo degrading into the SL clearly suggests that initiation of degradation also involves a mechanism that makes the 3’ end of the SL accessible to 3’hExo.

UPF1 likely plays a critical role in this initial degradation and our studies here suggest that it is recruited to the histone mRNP via interactions with the SLBP N-terminal region. We report that the N-terminal IDR of SLBP directly engages the helicase core of UPF1 to form a stable complex. The histone SL is not required for this interaction but facilitates it by stabilizing the SLBP protein in a well-folded conformation [47]. Since there is a requirement *in vivo* that the SL be close to the termination codon, the recruitment of UPF1 *in vivo* might also involve recognition of a ribosome bound to the termination site. The initial degradation steps occur on polyribosomes, and there is a clear barrier to degradation at the termination site, which is likely the ribosome [45].

Our findings are consistent with earlier observations of an RNA-independent interaction between UPF1 and SLBP [14]. The UPF1-SLBP interaction is unusual in that it is mediated by the UPF1 helicase core and does not involve the flanking CH or SQ domains that bind other factors involved in UPF1-mediated mRNA decay pathways, such as UPF2, STAU1, SMG5/7 and SMG6 [24, 32, 42, 48]. In addition to UPF1, the N-terminal IDR of SLBP engages factors involved in nuclear export and translation of histone mRNA, and harbors sites for phosphorylation by the Cyclin A/CDK1 complex [28, 49, 50]. It is not known if these interactions involve similar or distinct sites on the SLBP N-terminal IDR, which would lead to concomitant or mutually exclusive interactions being mediated by SLBP. Mutually exclusive interactions would ensure involvement of SLBP in only one histone mRNA processing step at a time. Multiple interactions with different partners involved in different processing steps would imply a spatial/temporal separation of the different steps in histone mRNA metabolism or a handover from one step to the next, allowing for efficient gene expression of the histone mRNA. Our biochemical analyses indicate that SLBP does not contain a specific binding site or motif to recognize UPF1 as successive truncations of the N-terminal IDR progressively weaken the UPF1-SLBP interaction. We envisage a binding mode where the SLBP-N-terminal IDR wraps around UPF1, making multiple contacts with the helicase core domains. Our crosslinking mass-spectrometry analysis are consistent with this model as UPF1-SLBP interlinks were mostly distributed over the RecA1, 1B and 1C domains of UPF1, with only a few interlinks to the UPF1-RecA2 domain (Figures 2A and 2B). Mapping of the crosslinked residues to the X-ray crystal structure of RNA-bound UPF1 show that the crosslinked residues cluster around the 3’-end of the RNA (Figure 2B), suggesting that binding of UPF1 to the single-stranded RNA stretch upstream of the SL positions it close to the SL RNA-bound SLBP (Figure 7C). Accordingly, interlinks were mapped to the SLBP-RBD even though this domain did not interact with UPF1 in our biochemical assays, consistent with an initial recognition event, which is followed by a more extensive interaction with the helicase domain..

The only other known example of a protein-protein interaction mediated by the UPF1 helicase core is an intra-molecular interaction with its own C-terminal SQ domain [32]. Binding of the SQ domain to the UPF1 helicase core inhibits its nucleic acid unwinding activity, possibly by restricting the conformational dynamics essential for a catalytic cycle. Similarly, we find that binding of SLBP to the UPF1 helicase domains hinders unwinding of nucleic acid duplexes. An increase in SLBP is accompanied by a concomitant decrease in UPF1 catalytic activity, suggesting a low affinity interaction or a complex with high kinetic off-rates. Presence of the SL does not hinder unwinding of the nucleic acid hybrid at the 3’-end. However, the SL-bound SLBP acts as an impediment for UPF1, evidenced by slowing down of unwinding activity in the early phase of the reaction. We suggest that UPF1 removes the impediment by displacing SLBP from the SL RNA and unwinding the SL, allowing it to translocate towards the 3’-end where it then unwinds the nucleic acid hybrid.

The binding and inhibition of UPF1 by SLBP is reminiscent of the inhibition of UPF1 by the poly-pyrimidine tract binding protein PTBP1, where binding of PTBP1 to UPF1 leads to dosage-dependent inhibition of UPF1 unwinding activity [33, 51]. However, the molecular mechanisms dictating the two events are very different. Binding of PTBP1 to UPF1 triggers dissociation of UPF1 from the target mRNA, protecting it from decay [33], whereas binding of SLBP to UPF1 recruits it to the histone mRNA to promote decay. This is evident from our observation that disruption of the UPF1-SLBP interaction slows down the rate of histone mRNA decay. This raises the obvious question: why does SLBP slow down UPF1 unwinding activity, if this activity is essential for decay. The answer to this, we argue, lies in achieving a balance between 5’-3’ unwinding of the SL by UPF1 and 3’-5’ degradation of the SL by 3’hExo, and ensuring that once recruited, UPF1 actively remodels the SLBP-bound SL upon receiving the correct cues for degradation (Figure 7C).

3’hExo is a member of the DEDDh family of exonucleases [13]. Accumulation of distinct intermediates in histone mRNA decay and *in vitro* studies suggests that it is a distributive enzyme that dissociates from its substrate after each round of catalysis [12, 36, 39, 40]. Atypical of DEDDh exonucleases is the presence of the SAP domain that confers RNA binding affinity and specificity for the histone SL [12]. The SAP domain recognizes the loop of the SL, rationalizing our observation that 3’hExo binds the intact SL and SL-decay intermediates with similar affinity (Figure 5A) [9]. While tethered to the SL (in the presence of SLBP) the 3’hExo cannot access the 3’ end of the histone mRNA to remove more than 2 or 3 nts, and the cell maintains this length of 3’ end via uridylation by TUT7 if additional nts are removed [34, 35]. Tethering of 3’hExo to the histone SL via its SAP domain presents two possible scenarios with respect to its ribonucleolytic activity, either 3’hExo dissociates from the RNA after removal of each nucleotide and rebinds the RNA using its SAP domain, which would be possible if the SAP domain has high on- and off-rates for the SL. Alternatively, the SAP domain remains bound to the loop of the SL while the nuclease domain dissociates from the substrate and is repositioned on the RNA for a new round of catalysis. This would require the SLBP binding to the SL to be perturbed. The 10-residue long flexible linker connecting the SAP and nuclease domains would allow 3’hExo to bind such that its nuclease domain can access the short single-stranded overhang at the 3’-end of the RNA [13].

Based on observations in previous reports and this study, we speculate that 3’hExo favours the second mode of sequential degradation. First, our *in vitro* degradation assay shows accumulation of distinct intermediates as opposed to a regular ladder consisting of intermediates differing by one nucleotide (Figure 5B). This is further supported by 3’-End-Seq analysis of histone mRNA decay intermediates in cells, where an uneven distribution of intermediates resulting from degradation into the stem of histone SL was observed [39]. Addition of a large excess of 3’hExo enhanced the speed of degradation but did not change the pattern of accumulation of intermediates. This suggests that the partially degraded pool of SL RNA is never fully accessible to the excess free 3’hExo in solution, which would degrade the RNA intermediates faster than they can be captured in such an assay. Likewise, unwinding of the SL by UPF1 increased the rate of degradation without affecting the degradation pattern of 3’hExo. We rationalize that UPF1 assists in repositioning of the 3’hExo nuclease domain on RNA by generating a small single-stranded overhang at the 3’-end.

Given the role of UPF1 in remodeling the SL-RNP and facilitating exoribonucleolytic decay, it is imperative that UPF1 catalytic activity is stringently regulated in this context. The catalytic activity of UPF1 is stimulated upon binding to UPF2, which also results in a weakening of the affinity of the helicase for RNA [20, 21, 23, 24]. This implies that recruitment of UPF2 to a UPF1-mediated decay pathway must either be transient or mediated through another factor. We find that the UPF2 binds 3’hExo with low affinity, using a sliding interaction motif that can similarly engage the SAP and nuclease domains of 3’hExo (Figure 6). The involvement of the acidic linker of UPF2 suggests that UPF2 binds the RNA binding groove of 3’hExo, which is lined with basic residues. The model of 3’hExo’s association with the histone SL via its SAP domain alone (in the “resting” state, when not actively degrading RNA) also allows us to imagine a state where the nuclease domain binds UPF2 instead of RNA. This would recruit UPF2 to the histone mRNP, which can then transiently activate UPF1 to unwind the SL. Following unwinding, the 3’hExo nuclease domain would release UPF2 to bind and degrade the RNA, before returning to its resting state (Figure 7C). The formation of a stable UPF2-3’hExo complex in solution (Figure 6D), despite the low affinity interaction and the formation of a stable UPF1-UPF2-3’hExo complex lend support to this model. Indeed, in every UPF1-mediated decay pathway investigated mechanistically thus far, UPF2 was shown to be anchored at the decay-RNP via a component other than UPF1. Interestingly, UPF1 is a far more abundant protein in cells and is present in ∼10-fold excess of UPF2 and 3’hExo [52], underscoring the importance of maintaining UPF2 near the helicase to ensure its rapid activation, particularly when timing of decay is crucial. In our previous siRNA experiments we did not see an effect of knocking down UPF2 on histone mRNA degradation [14], but this could be due to a transient requirement of UPF2. We also did not see an effect of efficient knockdown of 3’ hExo on histone mRNA degradation either, although the knockout prevented regulated histone mRNA degradation at the end of S-phase [34, 35].

The low speed of exoribonucleolytic decay by 3’hExo in solution as well as the accumulation of distinct decay intermediates in cells points to a slow initial decay phase. Notably, there is a clear absence of decay intermediates once 3’-5’ decay has progressed beyond the stem-loop. This can be attributed to recruitment of the bulk mRNA decay machinery (such as the cytoplasmic RNA exosome) to the histone mRNA [37]. It appears therefore that the role of 3’hExo is not only to initiate mRNA decay but also to prime the substrate for rapid decay with the assistance of other factors such as SLBP, UPF1, TUT7 and the LSm 1-7 complex [35, 53]. Together these factors ensure precision in timing of mRNA decay, both in initiation of the early slow decay phase and its transition to a later rapid decay phase. As such, the composition of the initial mRNP in histone mRNA decay bears remarkable similarity to a degradosome, a multi-protein assembly that contains an RNA helicase, a ribonuclease and a “sensor” that dictates the timing of target mRNA decay [54]. We propose that the sensor in histone mRNA decay could be an event in the lifetime of the histone mRNA, such as translation termination or recruitment of the additional factors mentioned above. Alternatively, modification of the proteins involved, such as dephosphorylation of SLBP which weakens its binding to the histone SL or phosphorylation of UPF1 by SMG1, which was shown to play a role in histone mRNA decay, could be the sensor that triggers decay [39, 55]. The complete details of how degradation is mechanistically achieved, particularly coupling of decay to translation termination on the histone mRNA, and the signal generated when DNA replication is inhibited in S-phase or at the S-G2 phase transition of the cell cycle remain questions for further studies.

## Methods

### Expression and purification of recombinant proteins from E. coli

The cDNAs of human 3’hExo, UPF1 and UPF2 and their respective truncations were cloned into a modified pET28a vector with an N-terminal 6x His-tag, followed by a TEV or HRV-3C protease cleavage site, using ligation independent cloning. For expression of GST-tagged proteins, a vector containing an N-terminal tandem 6x His-GST tag followed by a TEV or 3C protease cleavage site was used. Proteins were expressed in *Escherichia coli* BL21 Star(DE3) pRARE (3’hExo) or BL21 Gold(DE3) pLysS (UPF1 and UPF2). Cells were grown in terrific broth (TB) under antibiotic selection at 37 °C for 18 h or until an OD_600nm_ of ∼2.5 was reached. The temperature was reduced to 18 °C and expression was induced by addition of 0.1 mM IPTG (final concentration). Cells were harvested after 16-18 h and pelleted. Cell pellets were resuspended in lysis buffer (50 mM Tris-HCl pH 7.5, 500 mM/1 M NaCl, 10% glycerol, 1 mM MgCl_2_, 10 mM imidazole) and supplemented with DNase I and 1 mM PMSF. All buffers used for purification of UPF1-CHh were further supplemented with 1 µM ZnCl_2_. Cells were lysed by sonication and the lysate was clarified by centrifugation. The recombinant protein was enriched from the cell lysate using Ni^2+^-affinity chromatography. Protein bound Ni-NTA beads were extensively washed with lysis buffer. In the case of UPF1-CHh, an additional wash step with chaperone wash buffer (50 mM Tris-HCl pH 7.5, 1 M NaCl, 10% glycerol, 10 mM MgCl_2_, 50 mM KCl, 2 mM ATP, and 10 mM imidazole) was included. Following a final wash step with low salt buffer (50 mM Tris-HCl pH 7.5, 150 mM NaCl, 10% glycerol, 1 mM MgCl_2_ and 10 mM imidazole). 6x-His tagged proteins were eluted from the Ni-NTA beads with elution buffer (50 mM Tris-HCl pH 7.5, 150 mM NaCl, 10% glycerol, 1 mM MgCl_2_ and 300 mM imidazole). Proteins were further purified with a HiTrap Heparin Sepharose HP column (GE Healthcare) using heparin buffers A (20 mM Tris-HCl pH 7.5, 10% glycerol, 1 mM MgCl_2_, and 2 mM DTT) and B (20 mM Tris-HCl pH 7.5, 1 M NaCl, 10% glycerol, 1 mM MgCl_2_, and 2 mM DTT). Proteins were eluted from the column using a linear concentration gradient of NaCl and purified by a final size-exclusion chromatography step using Superdex 75 or Superdex 200 columns (Cytiva) in SEC buffer (3’hExo constructs: 20 mM HEPES pH 7.5, 150 mM NaCl, 5% glycerol, and 2 mM DTT; UPF1 and UPF2 constructs: 20 mM Tris-HCl pH 7.5, 150 mM NaCl, 5% glycerol, 1 mM MgCl_2_, and 2 mM DTT). Peak fractions were pooled, concentrated, flash frozen in liquid nitrogen and stored at −80°C until further use. The purity of the proteins after each chromatography step was assessed by SDS-PAGE analysis and Coomassie staining.

For NMR spectroscopy proteins were recombinantly labelled with ^15^N and/or ^13^C stable isotopes in M9 minimal medium. *E. coli* cells were transformed with the desired expression plasmid, grown to an OD_600_nm of ∼ 3.0 in TB media, spun down, and resuspended in an equal volume of minimal media. The cultures were then grown for 40 min at 37 °C to allow for recovery of the cells, followed by the addition of 8 g of glucose and 0.5 g of ^15^NH_4_Cl in the case of ^15^N-labelled proteins, or 2 g of ^13^C-glucose for ^13^C/^15^N labelling. The cultures were induced with 0.5 mM IPTG at 18 °C for 16-20 hours and centrifuged to collect the cell pellets. These were stored at -20°C until further processing. Purification of isotopically labelled proteins was performed as described above with no modifications. The protein samples were dialyzed against 20 mM HEPES (pH 7.0), 100 mM NaCl, 2 mM DTT, and concentrated to 100-500 μM using Amicon centrifugal filters (MWCO 3.5 kDa). The concentrations were determined using the absorbance at 280 nM and the predicted molar absorptivity based on the protein sequence [56].

### Expression and purification of SLBP from insect cells

Bacmids for insect cell transformation and recombinant virus expressing full-length human SLBP and SLBP variants were generated using the Bac-to-Bac™ baculovirus expression system. Baculoviruses were generated by transfection of the bacmid in Sf9 cells (cultured in Sf-900 II media) and further amplified by infection of Sf9 cells with a low-titer virus. For protein expression, Hi5 cells cultured in Express Five medium supplemented with glutamine were infected with 6xHis-TEV-SLBP viruses. Cells were harvested after 60 h by centrifugation at low speed. For purification, cells were resuspended in reduced salt buffer (10 mM Tris-HCl pH7.5, 10 mM NaCl) supplemented with DNase I and 1 mM PMSF. After mild sonication, the cell lysate was supplemented with 5x Nickel binding buffer such that a final buffer composition of 20 mM Tris-HCl pH 7.5, 400 mM NaCl, 10 mM Imidazole and 10 % glycerol was achieved. SLBP proteins were extracted from the lysate by Ni^2+^-affinity chromatography and further purified by Heparin-affinity chromatography, as described above. A final size-exclusion chromatography step was performed using a Superdex 75 column (Cytiva) in SEC buffer (20 mM Tris pH 7.5, 300 mM NaCl, 5% glycerol and 2 mM DTT). Peak fractions were pooled, concentrated, flash frozen in liquid nitrogen and stored at −80°C until further use. As with *E. coli* expressed proteins, an SDS-PAGE analysis was conducted after each step to monitor the purity of the proteins.

### GST pulldown assays

8 µg GST-tagged bait protein was incubated with 8 µg prey protein for 1 h, at 25 °C in GST-pulldown buffer (20 mM HEPES pH 7.5, 70 mM NaCl, 0.1 % NP-40 and 10 % glycerol). Samples were washed three times with GST-pulldown buffer before elution with GSH elution buffer (30 mM Tris pH 7.5, 150 mM NaCl, 1 mM MgCl_2_, 20 mM glutathione, 0.1 % NP-40, 14 % glycerol, 2 mM DTT buffer). Input and elution samples were resolved on an SDS PAGE gel and stained with Coomassie Brilliant Blue.

### Size exclusion chromatography (SEC)

For complex formation between UPF1 and SLBP, 1000 pmol of each protein and synthetic histone SL26 RNA (IDT), added in 1.2-fold molar excess wherever mentioned, were mixed to a final volume of 50 µl in analytical SEC buffer (20 mM HEPES pH 7.5, 75 mM NaCl, 5 % glycerol, 1 mM MgCl_2_, 2 mM DTT). For analysis of 3’hExo and UPF2 interactions, 1000 pmol of 3’hExo and 2000 pmol of UPF2-AL were used for the analysis. Samples were incubated overnight at 4 °C and resolved on a Superdex 200 Increase 3.2/300 column (GE Healthcare). The peak fractions were analyzed by SDS-PAGE, followed by staining with Coomassie brilliant blue. Protein-RNA complexes were additionally analysed on a 10 % urea-PAGE gel and stained with ethidium bromide.

The 3’hExo-UPF1-UPF2 complex was reconstituted on a preparative scale, using UPF1-CHh, UPF2_S_ and a variant of 3’hExo lacking the first 60 residues (3’hExoΔN). The proteins were mixed in equimolar amounts, incubated overnight at 4 °C and resolved on a Superdex 200 16/600 column using SEC buffer (20 mM HEPES pH 7.5, 100 mM NaCl, 2 % glycerol, 1 mM MgCl_2_, 2 mM DTT). The peak fractions were analysed by SDS-PAGE as above.

### Crosslinking mass spectrometry

To reconstitute the UPF1-SLBP-SLRNA complex on a preparative scale for CXMS, UPF1-Hel, SLBP-N and SL RNA were mixed at a molar ratio of 1.2:1:1.2, with the total protein amounting to ∼8 mg. The mixture was diluted to 1-2 mg/mL and incubated at room temperature for 30 min, followed by dialysis against CXMS-SEC buffer (20 mM Tris pH 7.5, 75 mM NaCl, 5 % Glycerol, 1 mM MgCl_2_ and 2 mM DTT) overnight at 4 °C. The mixture was concentrated and injected onto a Superdex 200 10/300 Increase column to separate the ternary complex from individual components and sub-complexes. The peak fractions were analysed by SDS-PAGE and Coomassie staining to detect proteins as well as urea-PAGE and ethidium bromide staining to detect RNA. Fractions corresponding to the ternary protein-RNA complex were carefully pooled and concentrated to 1.5 mg/mL for crosslinking.

For crosslinking, the complex was first exchanged into CX-buffer (20 mM HEPES, pH 7.5, 75 mM NaCl, 5 % Glycerol, 1 mM MgCl_2_). 10 µg aliquots of the protein complex were initially crosslinked with 0.2 mM, 0.5 mM, 1 mM and 2 mM BS3 or SMPB (Thermo Scientific) to optimize the minimum concentration of crosslinker necessary to achieve efficient crosslinking. A control reaction without crosslinker was performed in each case. The final crosslinking reaction was carried out with 1 mM BS3 (or 2 mM SMPB) for 30 minutes at room temperature and quenched with 50 mM Tris. Samples were separated by SDS-PAGE (NuPAGE 4-12 % gradient gel, Invitrogen). The crosslinked complex was cut out of the gel and separated into three pieces. Excised gel pieces were then subjected to in-gel tryptic digest. Samples were reduced with 10 mM dithiotreitol and alkylated with 55 mM iodacetamide and subsequently digested with trypsin (sequencing grade, Promega) at 37 °C for 18 hours. Extracted peptides were dried in a SpeedVac Concentrator and dissolved in loading buffer composed of 4 % acetonitrile and 0.05 % TFA. Samples were subjected to liquid chromatography mass spectrometry (LC-MS) on a QExactive HF-X (Thermo Scientific). Peptides were loaded onto a Dionex UltiMate 3000 UHPLC+ focused system (Thermo Scientific) equipped with an analytical column (75 µm x 300 mm, ReproSil-Pur 120 C18-AQ, 1.9 µm, Dr. Maisch GmbH, packed in house). Separation by reverse-phase chromatography was done on a 60-minute multi-step gradient with a flow rate of 0.3-0.4 µl min^-1^. MS1 spectra were recorded in profile mode with a resolution of 120 k, maximal injection time was set to 50 ms and AGC target to 1e^6^ to acquire a full MS scan between 380 and 1580 m/z. The top 30 abundant precursor ions (charge state 3-8) were triggered for HCD fragmentation (30 % NCE). MS2 spectra were recorded in profile mode with a resolution of 30 k; maximal injection time was set to 128 ms, AGC target to 2e^5^, isolation window to 1.4 m/z and dynamic exclusion was set to 30 seconds. Raw files were analyzed via pLink2.3.5 to identify crosslinked peptides, with standard settings changed as follows -peptide mass: 600-10000, precursor tolerance: 10 ppm, fixed modification: Carbamidomethyl [C] and variable modification Oxidation: [M] [57]. FDR was set to 1 % and results were filtered by excluding crosslinks supported by only one crosslinked peptide spectrum match. The interaction network for the ternary complex was illustrated via xiNET [58]. The analysis was carried out with one sample (three technical replicates for each crosslinker).

### CRISPR/Cas9 mediated SLBP exon 2 deletion and RT-PCR analysis

SLBP exon 2 was deleted in HCT116 cells using CRISPR/Cas9 as described in [34]. Four sgRNAs targeting the first and second introns flanking exon 2 were designed using CRISPOR (http://crispor.tefor.net/). These sgRNAs were then synthesized with ‘ccgg’ and ‘aaac’ modifications at the 5’ ends of the forward and reverse strands, respectively, to match the sticky ends of the vector after annealing. The sgRNAs were phosphorylated and ligated into a BsaI-digested sgRNA vector (Addgene #51133). HCT116 cells were cultured in McCoy’s 5A medium (Corning, 10-050-CV) and approximately 0.4 x 10^6^ cells were seeded in six-well plates. Transfection was performed 24 hours later. The correctly sequenced sgRNA plasmids were co-transfected with Cas9 (Addgene #44758) and GFP (pmaxGFP from Lonza) plasmids into cells using jetOPTIMUS (Genesee Scientific). GFP positive single cells were isolated with the CellRaft Air® System and cultured in 96-well plates. Genomic PCR, with primers designed to amplify a 500 bp region around the target sites, was used to confirm the presence of the mutation. Positive PCR products were sequenced (Eton Bioscience) and cloned into a pJET vector (Thermo Fisher, K1231) for allelic sequence determination.

To confirm exon 2 deletion at the mRNA level, RT-PCR was performed. RNA was extracted from cells and cDNA was synthesized using a reverse transcriptase kit (Applied Biostems 4387406). PCR amplification was carried out with a forward primer located in exon 1 and a reverse primer spanning the exon 4-exon 5 junction. This strategy ensured that only the mRNA-transcribed cDNA was amplified, while any genomic DNA contamination was eliminated by the primer design. The PCR products were analyzed by agarose gel electrophoresis to verify the absence of exon 2 in SLBPΔ2 cells.

### Western Blot

To verify exon 2 deletion at the protein level, western blotting was performed as described by Holmquist and co-workers [34]. Cells were collected and lysed using a buffer containing 0.5% NP-40, 50 mM Tris-HCl, 300 mM NaCl, and protease inhibitors (Roche, 11836170001). The lysates were incubated on ice for 20 minutes, followed by centrifugation at 4°C. Protein concentration was determined using a BCA protein assay. A total of 10 μg of protein was separated on a 4-12% gradient gel and transferred to a nitrocellulose membrane (Thermo REF 88018) at 100V for 1 hour. The membrane was blocked in 5% non-fat milk in phosphate buffered saline supplemented with 0.1 % Tween-20 for 1 hour and incubated overnight at 4°C with primary antibody in PBST. After washing, the membrane was incubated with a secondary antibody for 1 hour at room temperature. The membrane was washed again, and protein signals were developed using ECL for 3 minutes. Protein expression was visualized using a KwikQuant imaging system.

### Northern Blot

For study of histone mRNA decay, WT and SLBPΔE2 cells were treated with 5 mM HU (hydroxyurea, Sigma) for 20 and 40 minutes. Total RNA was extracted from untreated and HU-treated cells using Trizol reagent (Ambion, 15596026). RNA was quantified using a Nanodrop spectrophotometer. For Northern blotting, 5 μg of RNA was denatured by incubation at 95°C for 5 minutes and then separated by electrophoresis on a 6% AccuGel. RNA was transferred to a membrane (Cytiva RPN303B) using cold 0.5x TBE at 30V for 40 minutes. The membrane was dried for 1 hour and cross-linked with UV light at 1200 μJOULES x100 using a Stratalinker 2400 system. For hybridization, the membrane was prehybridized in hybridization buffer (Invitrogen, AM8670) at 42°C for 30 minutes. Probes were synthesized from purified PCR templates using random priming, denatured at 95°C, and hybridized to the membrane overnight at 42°C. The membrane was washed twice with 2% SSC and once with 0.01% SSC before being scanned for signals using a Typhoon imaging system.

### In vitro transcription of RNA substrates

For precise RNA 3’ ends, histone stem-loop RNAs were *in vitro* transcribed from a DNA template where the HDV-ribozyme sequence was fused to the 3’ end of the template for SL-RNA (see Supplementary figure 4 and Supplementary table 1). DNA template sequences were cloned into a pJet1.2/blunt cloning vector using the CloneJET PCR-cloning kit (ThermoFisher). PCR-amplified double-stranded (ds) DNA template was incubated with T7 RNA polymerase (purified in-house) together with 16 mM NTPs, 50 mM DTT, 20 mM MgCl_2_, 2 mM Spermidine, 6.4 % PEG 8000 and pyrophosphatase enzyme (purified in-house) in a 200 mM Tris-HCl pH 8 buffer for 4-6 h at 37 °C. The reaction was quenched with EDTA prior to gel purification of the transcribed RNA. The reaction mix consisting of different RNA fragments was resolved on a 10 % denaturing PAGE, from which the gel fragment corresponding to the auto-cleaved SL RNA was excised. Gel pieces were crushed and soaked overnight in RNA elution buffer (20 mM Tris-HCl, 300 mM sodium acetate, 2 mM EDTA). Recovered RNA in solution was buffer-exchanged into nuclease-free water (HyPure water, Cytiva) and concentrated using a 3-kDa molecular weight cut-off concentrator (Merck). To remove residual buffer components, RNA was precipitated out of solution using ethanol and redissolved in nuclease-free HyPure water. The purified RNA was stored at -20 °C until further use.

### Unwinding assay

Unwinding assays were based on those described by Fritz and co-workers [33] and were adapted for our purposes. RNA substrates were in vitro transcribed from PCR-amplified dsDNA templates, following the protocol above. Substrates for unwinding assays were not transcribed as precursors fused to HDV-ribozyme. 5’-Alexa Fluor488-labelled DNA that was used to create the RNA:DNA duplex and Blackhole Quencher 1 (BHQ1)-labeled DNA (used as a trap) was purchased from Integrated DNA Technologies (IDT). Sequences of all nucleic acids used in this experiment (including the DNA template for IVT) are provided in Supplementary table 1. For duplex annealing, the respective RNA and fluorescent DNA were mixed in a 11:7 ratio to a final concentration of 864 nM RNA and 543 nM DNA together with 2 mM magnesium acetate in 1x unwinding buffer (50 mM MES pH 6.5, 50 mM potassium acetate, 0.1 % NP-40). The RNA:DNA duplex was incubated at 96 °C for 4 min, snap cooled on ice for 30 min and used the same day. For each replicate, a total reaction volume of 40 μl was prepared with 1x unwinding buffer supplemented with 2 mM magnesium acetate and 2 mM DTT. 7.5 pmol of DNA:RNA duplex were mixed with 12 pmol of UPF1-Hel or UPF1CH-h protein, and wherever mentioned with12 or 20 pmol of the respective SLBP protein and/or 24 pmol UPF2_S_ in a volume of 22 µl. The reaction was incubated for 30 min at 25 °C. 35 pmol of BHQ1 DNA quencher was added, following which the reaction was transferred to a black, flat-bottomed, 384-well plate (Corning). Negative controls contained 1x unwinding buffer instead of protein or RNA. To start the reaction, 16 µl of 2 mM ATP in 1x unwinding buffer was added to each sample using the injector module of a Spark multimode microplate reader (Tecan). Fluorescence was monitored over a period of 60 min in 10 s intervals. Two technical duplicates were performed for each condition. The values were corrected for baseline fluorescence by subtracting the initial fluorescence reading immediately following addition of ATP. This was then normalized to the baseline-corrected maximum fluorescence value for each data set.

### Fluorescence Anisotropy assay

To determine the binding affinities of 3’hExo and SLBP to SL-RNA and SL degradation intermediates by fluorescence anisotropy assay, synthetic RNAs labelled with 6-FAM at their 5’-end were procured from IDT (see Supplementary Table 1 for RNA sequences). The respective RNA was incubated with increasing amounts of SLBPfl or 3’hExofl. A 45 µl binding-reaction containing RNA and protein was prepared in 1x fluorescence anisotropy buffer (20 mM HEPES pH 7.5, 100 mM NaCl, 1 mM MgCl_2_, 0.5 mM EDTA, supplemented with BSA to a final concentration of 100 µg/ml). The reactions were incubated for 30 min at room temperature. 40 μL of each sample was transferred to a black, flat-bottomed, 384-well plate (Corning). The negative controls were prepared with 1x fluorescence anisotropy buffer in place of the protein. Fluorescence polarization was measured with a multi-modal Tecan plate reader (Spark) at 25 °C. Technical duplicates were prepared and measured for each protein concentration. The values were corrected for background by subtracting the value obtained for the negative control lacking protein. Fluorescence anisotropy was calculated according to the formula 2·FP/(3-FP) and subsequently normalized to the maximum anisotropy value. Values were averaged and fitted to an equation representing one site-specific binding with a Hill slope using Prism software (GraphPad).

### Degradation assay

To monitor the degradation of SL RNA by 3’hExo, a time course experiment was performed using radiolabelled RNA, where the degradation products of the RNA were resolved on a denaturing PAGE. 0.1 nM radiolabelled RNA was incubated with 200 nM 3’hExofl (or 400 nM 3’hExofl for 2x 3’hExo), 200 nM cold RNA and where indicated, with 200 nM UPF1-Hel, in 1x degradation buffer (20 mM HEPES pH 7.5, 50 mM NaCl, 5 % glycerol, 2 mM DTT) on ice at 4 °C. The degradation reaction was initiated by addition of 5 mM MgCl_2_ and 1.7 mM ATP (final concentrations) and conducted at 25 °C. 10 µl were removed from the reaction mixture at indicated time points and were quenched with an equal volume of 2x RNA dye. For the 0 min time point, 10 µl was removed prior to the addition of MgCl_2_ and ATP. Degradation products were resolved on a 15 % denaturing PAGE. The RNA was visualized via phosphorimaging as described above for EMSAs.

### NMR spectroscopy

NMR experiments were collected on a Bruker Avance 600, 700, and 800 MHz spectrometers at 298 K (25 °C). The NMR samples contained 10 % D_2_O as a lock agent and 0.2 % sodium azide for increased sample stability. Titration experiments were performed using isotopically ^15^N-labelled UPF2^1015-1106^ or UPF2^1024-1085^, the latter used for K_D_ determination (see below). The backbone assignments of UPF2^1015-1106^ (^13^C^α^, ^13^C^β^, ^15^N, and ^1^H^N^) were obtained with ^15^N/^13^C-labeled samples and standard ^1^H-^13^C-^15^N scalar correlation experiments with apodization weighted sampling [59, 60]. The secondary structure propensity obtained from NMR chemical shifts was determined using the MICS algorithm [61].

To monitor changes occurring in UPF2 upon addition of 3’hExo, ^15^N-HSQC spectra were recorded after stepwise addition of unlabeled proteins, either full-length or containing the SAP (63-124) or nuclease (124-349) domains of 3’hExo. The resulting spectra were analyzed using NMRPipe and NMRFAM-Sparky [62, 63]. The CSP values in the presence of varying amounts of 3’hExo were calculated as Δδ = [(0.14Δδ_N_)^2^+ (Δδ_H_)^2^]^1/2^, where Δδ_N_ and Δδ_H_ are the changes in chemical shift for ^15^N and ^1^H^N^, respectively. To determine the K_D_, twelve well-resolved amide peaks exhibiting fast-exchange behavior upon addition of 3.75 molar excess of 3’hExo^FL^ to UPF2^1024-1085^ were tracked. The CSP values were plotted as a function of added 3’hExo^FL^, and the resulting titration curves fit with GraphPad Prism to the equation for a one-to-one binding isotherm as described previously [64]. The resulting mean value and standard deviation is reported.

## Author Contributions

A.M.A, G.X., T.D., S.L. and S.C.: protein purification, biochemical and biophysical experiments; W.H. and W.F.M.: analysis of histone mRNA decay in wild type and CRISPR/Cas9 edited cells; C.P-B. and J.H.: NMR spectroscopy and data analysis. J.B. and U.H.: crosslinking mass-spectrometry and data analysis; N.V., V.N. and C.K.: additional biochemical and biophysical experiments. A.M.A., W.F.M. and S.C.: experimental design and manuscript preparation; all authors: manuscript editing.

## Acknowledgements

We thank Rhese D. Thompson and Qi Zhang for sharing their expertise on *in vitro* transcription, Lea S. Pommerening for help with establishing nucleic acid unwinding assays, Florian Heyd for critically reading the manuscript and members of the Chakrabarti lab for helpful discussions. We gratefully acknowledge the BioSupraMol core facility at Freie Universität Berlin for access to the Tecan Spark plate reader and Biacore^TM^ X100. S.C., J.H. and H.U. are supported by grants from the Deutsche Forschungsgemeinschaft (DFG) and acknowledge funding through the priority program SPP1935. S.C. is additionally supported by the Heisenberg program of the DFG. C.P.-B. was supported by the EMBL Interdisciplinary Postdoc (EI_3_POD) Program fellowship under Marie Sklodowska-Curie Actions COFUND (grant no. 664726). W.F.M is supported by grants from the NIH.

## Data Availability

All original data and material will be made available upon reasonable request.

## References

1. Harris, M.E., et al., Regulation of histone mRNA in the unperturbed cell cycle: evidence suggesting control at two posttranscriptional steps. Mol Cell Biol, 1991. 11(5): p. 2416–24.

2. Heintz, N., H.L. Sive, and R.G. Roeder, Regulation of human histone gene expression: kinetics of accumulation and changes in the rate of synthesis and in the half-lives of individual histone mRNAs during the HeLa cell cycle. Mol Cell Biol, 1983. 3(4): p. 539–50.

3. Armstrong, C. and S.L. Spencer, Replication-dependent histone biosynthesis is coupled to cell-cycle commitment. Proc Natl Acad Sci U S A, 2021. 118(31).

4. Marzluff, W.F., Histone 3’ ends: essential and regulatory functions. Gene Expr, 1992. 2(2): p. 93–7.

5. Birnstiel, M.L., M. Busslinger, and K. Strub, Transcription termination and 3’ processing: the end is in site! Cell, 1985. 41(2): p. 349-59.

6. Wang, Z.F., et al., The protein that binds the 3’ end of histone mRNA: a novel RNA-binding protein required for histone pre-mRNA processing. Genes Dev, 1996. 10(23): p. 3028–40.

7. Dominski, Z., et al., Stem-loop binding protein facilitates 3’-end formation by stabilizing U7 snRNP binding to histone pre-mRNA. Mol Cell Biol, 1999. 19(5): p. 3561–70.

8. Battle, D.J. and J.A. Doudna, The stem-loop binding protein forms a highly stable and specific complex with the 3’ stem-loop of histone mRNAs. RNA, 2001. 7(1): p. 123–32.

9. Tan, D., et al., Structure of histone mRNA stem-loop, human stem-loop binding protein, and 3’hExo ternary complex. Science, 2013. 339(6117): p. 318-21.

10. Marzluff, W.F. and K.P. Koreski, Birth and Death of Histone mRNAs. Trends Genet, 2017. 33(10): p. 745–759.

11. Dominski, Z., et al., A 3’ exonuclease that specifically interacts with the 3’ end of histone mRNA. Mol Cell, 2003. 12(2): p. 295–305.

12. Yang, X.C., et al., Characterization of 3’hExo, a 3’ exonuclease specifically interacting with the 3’ end of histone mRNA. J Biol Chem, 2006. 281(41): p. 30447–54.

13. Cheng, Y. and D.J. Patel, Crystallographic structure of the nuclease domain of 3’hExo, a DEDDh family member, bound to rAMP. J Mol Biol, 2004. 343(2): p. 305–12.

14. Kaygun, H. and W.F. Marzluff, Regulated degradation of replication-dependent histone mRNAs requires both ATR and Upf1. Nat Struct Mol Biol, 2005. 12(9): p. 794–800.

15. Brooks, L., 3rd, et al., A multiprotein occupancy map of the mRNP on the 3’ end of histone mRNAs. RNA, 2015. 21(11): p. 1943-65.

16. Lavysh, D. and G. Neu-Yilik, UPF1-Mediated RNA Decay-Danse Macabre in a Cloud. Biomolecules, 2020. 10(7).

17. Kim, Y.K. and L.E. Maquat, UPFront and center in RNA decay: UPF1 in nonsense-mediated mRNA decay and beyond. RNA, 2019. 25(4): p. 407–422.

18. Franks, T.M., G. Singh, and J. Lykke-Andersen, Upf1 ATPase-dependent mRNP disassembly is required for completion of nonsense-mediated mRNA decay. Cell, 2010. 143(6): p. 938–50.

19. Chapman, J.H., et al., UPF1 mutants with intact ATPase but deficient helicase activities promote efficient nonsense-mediated mRNA decay. Nucleic Acids Res, 2022. 50(20): p. 11876–11894.

20. Chakrabarti, S., et al., Molecular mechanisms for the RNA-dependent ATPase activity of Upf1 and its regulation by Upf2. Mol Cell, 2011. 41(6): p. 693–703.

21. Clerici, M., et al., Unusual bipartite mode of interaction between the nonsense-mediated decay factors, UPF1 and UPF2. EMBO J, 2009. 28(15): p. 2293-306.

22. Kashima, I., et al., Binding of a novel SMG-1-Upf1-eRF1-eRF3 complex (SURF) to the exon junction complex triggers Upf1 phosphorylation and nonsense-mediated mRNA decay. Genes Dev, 2006. 20(3): p. 355–67.

23. Xue, G., et al., Modulation of RNA-binding properties of the RNA helicase UPF1 by its activator UPF2. RNA, 2023. 29(2): p. 178–187.

24. Chamieh, H., et al., NMD factors UPF2 and UPF3 bridge UPF1 to the exon junction complex and stimulate its RNA helicase activity. Nat Struct Mol Biol, 2008. 15(1): p. 85–93.

25. Gehring, N.H., et al., Exon-junction complex components specify distinct routes of nonsense-mediated mRNA decay with differential cofactor requirements. Mol Cell, 2005. 20(1): p. 65–75.

26. Melero, R., et al., The cryo-EM structure of the UPF-EJC complex shows UPF1 poised toward the RNA 3’ end. Nat Struct Mol Biol, 2012. 19(5): p. 498–505, S1-2.

27. Yamashita, A., et al., Human SMG-1, a novel phosphatidylinositol 3-kinase-related protein kinase, associates with components of the mRNA surveillance complex and is involved in the regulation of nonsense-mediated mRNA decay. Genes Dev, 2001. 15(17): p. 2215–28.

28. Zheng, L., et al., Phosphorylation of stem-loop binding protein (SLBP) on two threonines triggers degradation of SLBP, the sole cell cycle-regulated factor required for regulation of histone mRNA processing, at the end of S phase. Mol Cell Biol, 2003. 23(5): p. 1590–601.

29. Whitfield, M.L., et al., Stem-loop binding protein, the protein that binds the 3’ end of histone mRNA, is cell cycle regulated by both translational and posttranslational mechanisms. Mol Cell Biol, 2000. 20(12): p. 4188–98.

30. Sun, Y., et al., Structure of an active human histone pre-mRNA 3’-end processing machinery. Science, 2020. 367(6478): p. 700-703.

31. Fairman-Williams, M.E., U.P. Guenther, and E. Jankowsky, SF1 and SF2 helicases: family matters. Curr Opin Struct Biol, 2010. 20(3): p. 313–24.

32. Fiorini, F., M. Boudvillain, and H. Le Hir, Tight intramolecular regulation of the human Upf1 helicase by its N- and C-terminal domains. Nucleic Acids Res, 2013. 41(4): p. 2404–15.

33. Fritz, S.E., et al., The RNA-binding protein PTBP1 promotes ATPase-dependent dissociation of the RNA helicase UPF1 to protect transcripts from nonsense-mediated mRNA decay. J Biol Chem, 2020. 295(33): p. 11613–11625.

34. Holmquist, C.E., et al., Knockouts of TUT7 and 3’hExo show that they cooperate in histone mRNA maintenance and degradation. RNA, 2022. 28(11): p. 1519–1533.

35. Lackey, P.E., J.D. Welch, and W.F. Marzluff, TUT7 catalyzes the uridylation of the 3’ end for rapid degradation of histone mRNA. RNA, 2016. 22(11): p. 1673–1688.

36. Hoefig, K.P., et al., Eri1 degrades the stem-loop of oligouridylated histone mRNAs to induce replication-dependent decay. Nat Struct Mol Biol, 2013. 20(1): p. 73–81.

37. Mullen, T.E. and W.F. Marzluff, Degradation of histone mRNA requires oligouridylation followed by decapping and simultaneous degradation of the mRNA both 5’ to 3’ and 3’ to 5’. Genes Dev, 2008. 22(1): p. 50–65.

38. Holmquist, C.E. and W.F. Marzluff, Determining degradation intermediates and the pathway of 3’ to 5’ degradation of histone mRNA using high-throughput sequencing. Methods, 2019. 155: p. 104–115.

39. Meaux, S.A., C.E. Holmquist, and W.F. Marzluff, Role of oligouridylation in normal metabolism and regulated degradation of mammalian histone mRNAs. Philos Trans R Soc Lond B Biol Sci, 2018. 373(1762).

40. Slevin, M.K., et al., Deep sequencing shows multiple oligouridylations are required for 3’ to 5’ degradation of histone mRNAs on polyribosomes. Mol Cell, 2014. 53(6): p. 1020–30.

41. Clerici, M., et al., Structural and functional analysis of the three MIF4G domains of nonsense-mediated decay factor UPF2. Nucleic Acids Res, 2014. 42(4): p. 2673–86.

42. Gowravaram, M., et al., Insights into the assembly and architecture of a Staufen-mediated mRNA decay (SMD)-competent mRNP. Nat Commun, 2019. 10(1): p. 5054.

43. Kadlec, J., E. Izaurralde, and S. Cusack, The structural basis for the interaction between nonsense-mediated mRNA decay factors UPF2 and UPF3. Nat Struct Mol Biol, 2004. 11(4): p. 330–7.

44. Lopez-Perrote, A., et al., Human nonsense-mediated mRNA decay factor UPF2 interacts directly with eRF3 and the SURF complex. Nucleic Acids Res, 2016. 44(4): p. 1909–23.

45. Kaygun, H. and W.F. Marzluff, Translation termination is involved in histone mRNA degradation when DNA replication is inhibited. Mol Cell Biol, 2005. 25(16): p. 6879–88.

46. Ivanov, P.V., et al., Interactions between UPF1, eRFs, PABP and the exon junction complex suggest an integrated model for mammalian NMD pathways. EMBO J, 2008. 27(5): p. 736–47.

47. Zhang, J., et al., Molecular mechanisms for the regulation of histone mRNA stem-loop-binding protein by phosphorylation. Proc Natl Acad Sci U S A, 2014. 111(29): p. E2937–46.

48. Okada-Katsuhata, Y., et al., N- and C-terminal Upf1 phosphorylations create binding platforms for SMG-6 and SMG-5:SMG-7 during NMD. Nucleic Acids Res, 2012. 40(3): p. 1251–66.

49. Erkmann, J.A., et al., Nuclear import of the stem-loop binding protein and localization during the cell cycle. Mol Biol Cell, 2005. 16(6): p. 2960–71.

50. von Moeller, H., C. Basquin, and E. Conti, The mRNA export protein DBP5 binds RNA and the cytoplasmic nucleoporin NUP214 in a mutually exclusive manner. Nat Struct Mol Biol, 2009. 16(3): p. 247–54.

51. Fritz, S.E., et al., An alternative UPF1 isoform drives conditional remodeling of nonsense-mediated mRNA decay. EMBO J, 2022. 41(10): p. e108898.

52. Cho, N.H., et al., OpenCell: Endogenous tagging for the cartography of human cellular organization. Science, 2022. 375(6585): p. eabi6983.

53. Lyons, S.M., et al., The C-terminal extension of Lsm4 interacts directly with the 3’ end of the histone mRNP and is required for efficient histone mRNA degradation. RNA, 2014. 20(1): p. 88–102.

54. Machado de Amorim, A. and S. Chakrabarti, Assembly of multicomponent machines in RNA metabolism: A common theme in mRNA decay pathways. Wiley Interdiscip Rev RNA, 2021: p. e1684.

55. Krishnan, N., et al., The prolyl isomerase Pin1 targets stem-loop binding protein (SLBP) to dissociate the SLBP-histone mRNA complex linking histone mRNA decay with SLBP ubiquitination. Mol Cell Biol, 2012. 32(21): p. 4306–22.

56. Gasteiger, E., et al., ExPASy: The proteomics server for in-depth protein knowledge and analysis. Nucleic Acids Res, 2003. 31(13): p. 3784–8.

57. Chen, Z.L., et al., A high-speed search engine pLink 2 with systematic evaluation for proteome-scale identification of cross-linked peptides. Nat Commun, 2019. 10(1): p. 3404.

58. Combe, C.W., L. Fischer, and J. Rappsilber, xiNET: cross-link network maps with residue resolution. Mol Cell Proteomics, 2015. 14(4): p. 1137–47.

59. Simon, B. and H. Kostler, Improving the sensitivity of FT-NMR spectroscopy by apodization weighted sampling. J Biomol NMR, 2019. 73(3-4): p. 155–165.

60. Sattler, M., Schleucher, J., Griesinger, C., Heteronuclear multidimensional NMR experiments for the structure determination of proteins in solution employing pulsed field gradients. Progress in Nuclear Magnetic Resonance Spectroscopy, 1999. 34(2): p. 93–158.

61. Shen, Y. and A. Bax, Identification of helix capping and b-turn motifs from NMR chemical shifts. J Biomol NMR, 2012. 52(3): p. 211–32.

62. Lee, W., M. Tonelli, and J.L. Markley, NMRFAM-SPARKY: enhanced software for biomolecular NMR spectroscopy. Bioinformatics, 2015. 31(8): p. 1325–7.

63. Delaglio, F., et al., NMRPipe: a multidimensional spectral processing system based on UNIX pipes. J Biomol NMR, 1995. 6(3): p. 277–93.

64. Perez-Borrajero, C., et al., The Biophysical Basis for Phosphorylation-Enhanced DNA-Binding Autoinhibition of the ETS1 Transcription Factor. J Mol Biol, 2019. 431(3): p. 593–614.

