## Supplementary figure for "Molecular mechanisms of recruitment, function and regulation of UPF1 in histone mRNA decay"

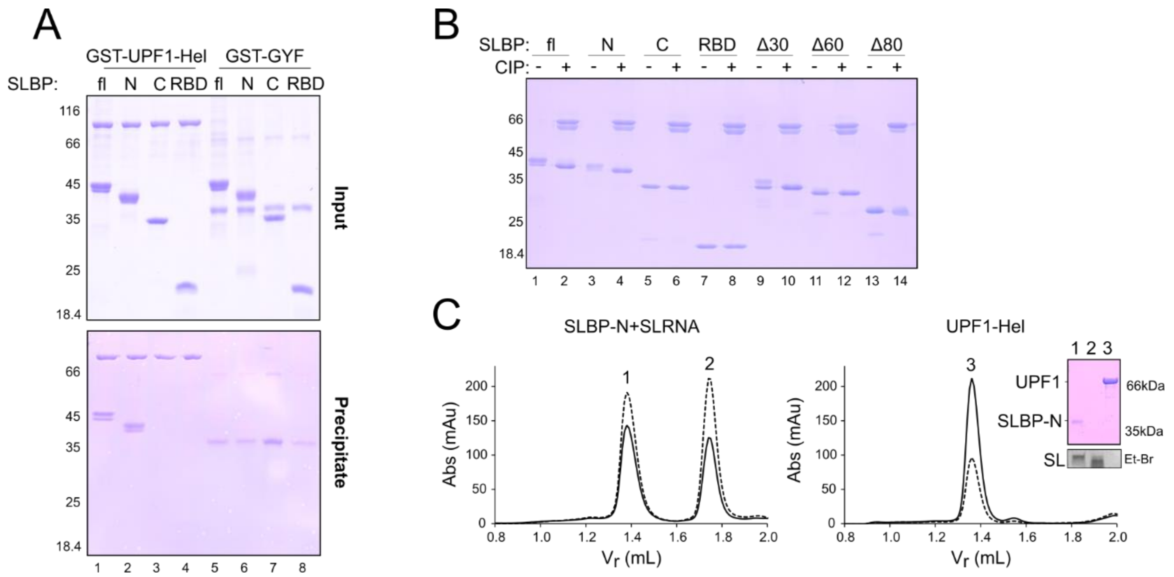


**Supplementary figure 1** (related to Figure 1)

**(A)** The complete SDS-PAGE gel corresponding to Figure 1C, including negative controls using GST-GYF as a bait. SLBP variants used in this experiment are shown in Figure 1A.

**(B)** SDS-PAGE analysis of phosphatase-treatment of SLBP protein. Equal amounts (2 μg) of each SLBP variant were treated with 5 units of calf intestinal phosphatase (CIP) for 30 minutes at 30 °C, following which the reaction was quenched by addition of 4X SDS-PAGE buffer. Mock-treated (-) and CIP-treated (+) samples were analysed by SDS-PAGE. Compaction of the bands corresponding to SLBPfl, SLBP-N, SLBPΔ30 and SLBPΔ60 and the concomitant increase in mobility on SDS-PAGE suggests that these proteins were phosphorylated upon expression in insect cells. SLBP-C and RBD did not show any change in migration pattern on SDS-PAGE upon CIP treatment.

**(C)** Analytical size exclusion chromatography (SEC) of an SLBP-N:SL26 complex (left panel) and UPF1-Hel (middle panel). The corresponding SDS- and urea-PAGE analyses of the peak fractions are shown on the right. Appearance of a lower retention volume peak (1.25 mL) upon addition of SL26 RNA to the UPF1:SLBP protein mixture (Figure 1D) indicates formation of a stable UPF1:SLBP complex only when SLBP is bound to the histone SL RNA.

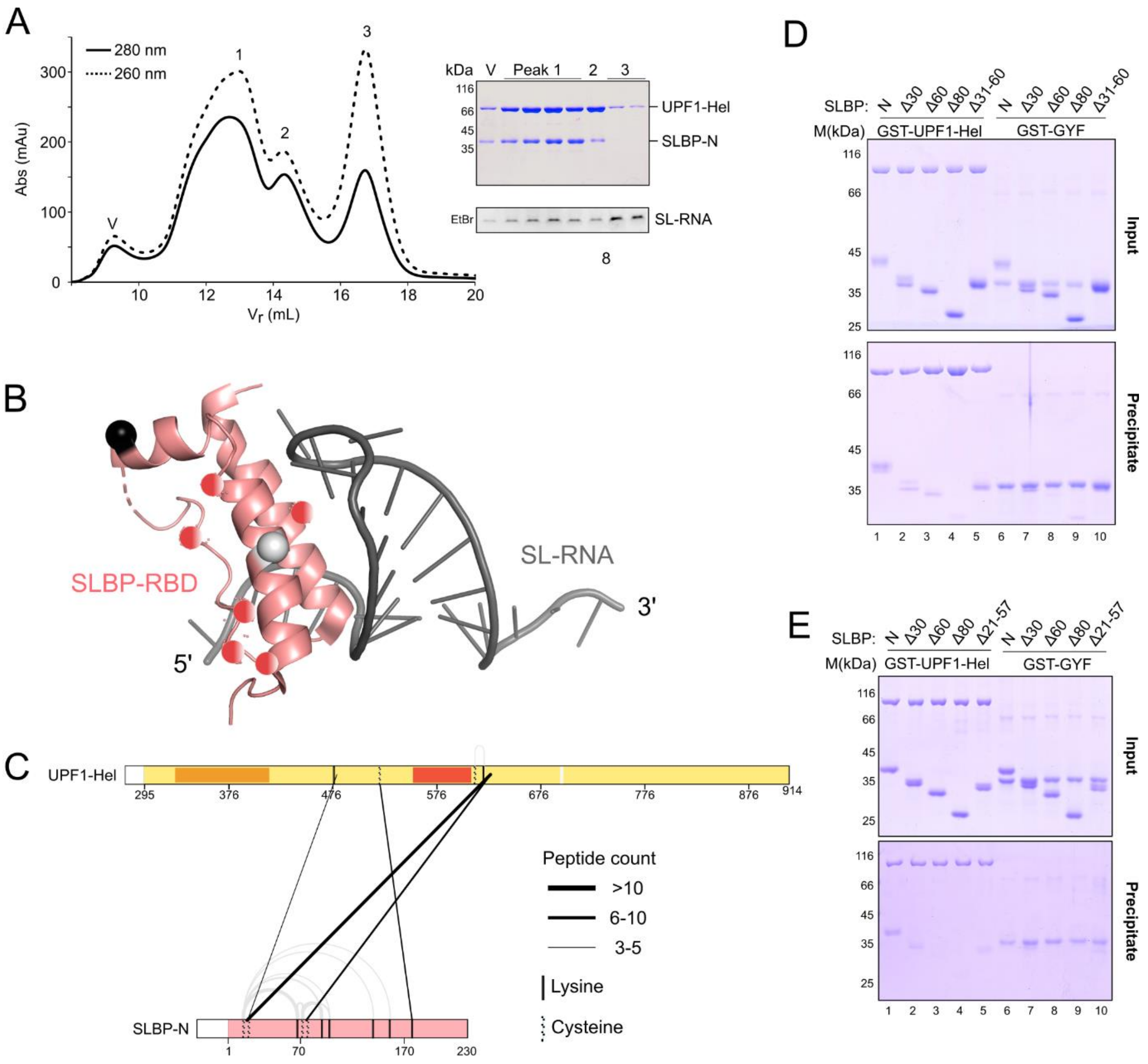


**Supplementary figure 2** (related to figure 2)

**(A)** Preparative SEC for the reconstitution of a ternary complex of UPF1-Hel, SLBP-N and U12-SL26 RNA for crosslinking mass-spectrometry (CLMS). The chromatogram of the SEC run and the corresponding SDS- and urea-PAGE analyses are shown on the left and right, respectively. Peak fractions of peak 1 were pooled for CLMS analysis.

**(B)** X-ray crystal structure of SLBP/SL26-RNA (derived from the X-ray crystal structure of the SLBP/3'hExo/SL26 complex, PDB 4L8R), with the position of the residues within the SLBP-RBD that crosslink to UPF1 (using BS3) highlighted as spheres. The color scheme of the spheres (black, and light grey) corresponds to the color of the lines denoting interlinks in (C). Lysine residues within the SLBP-RBD that were not crosslinked to UPF1 are shown as pink spheres.

**(C)** Linkage map showing the interlinks between UPF1 and SLBP derived from CLMS analysis of the UPF1-Hel:SLBP-N:12U-SL26 RNA complex, using the crosslinker SMPB. Intra- and Interlinks are depicted as in Figure 2A. Fewer crosslinks were obtained with SMPB in comparison to BS3 (refer to Figure 2A). All interlinks between SLBP and UPF1 mapped to the RecA1 domain of UPF1.

**(D)** The complete SDS-PAGE gel corresponding to Figure 2C, including negative controls using GST-GYF as a bait. SLBP variants used in this experiment are shown in Figure 1A.

**(E)** GST-pulldown assay of GST-UPF1-Hel (and GST-GYF) with SLBP N-terminal truncations including SLBP$\Delta$31-60, as above.

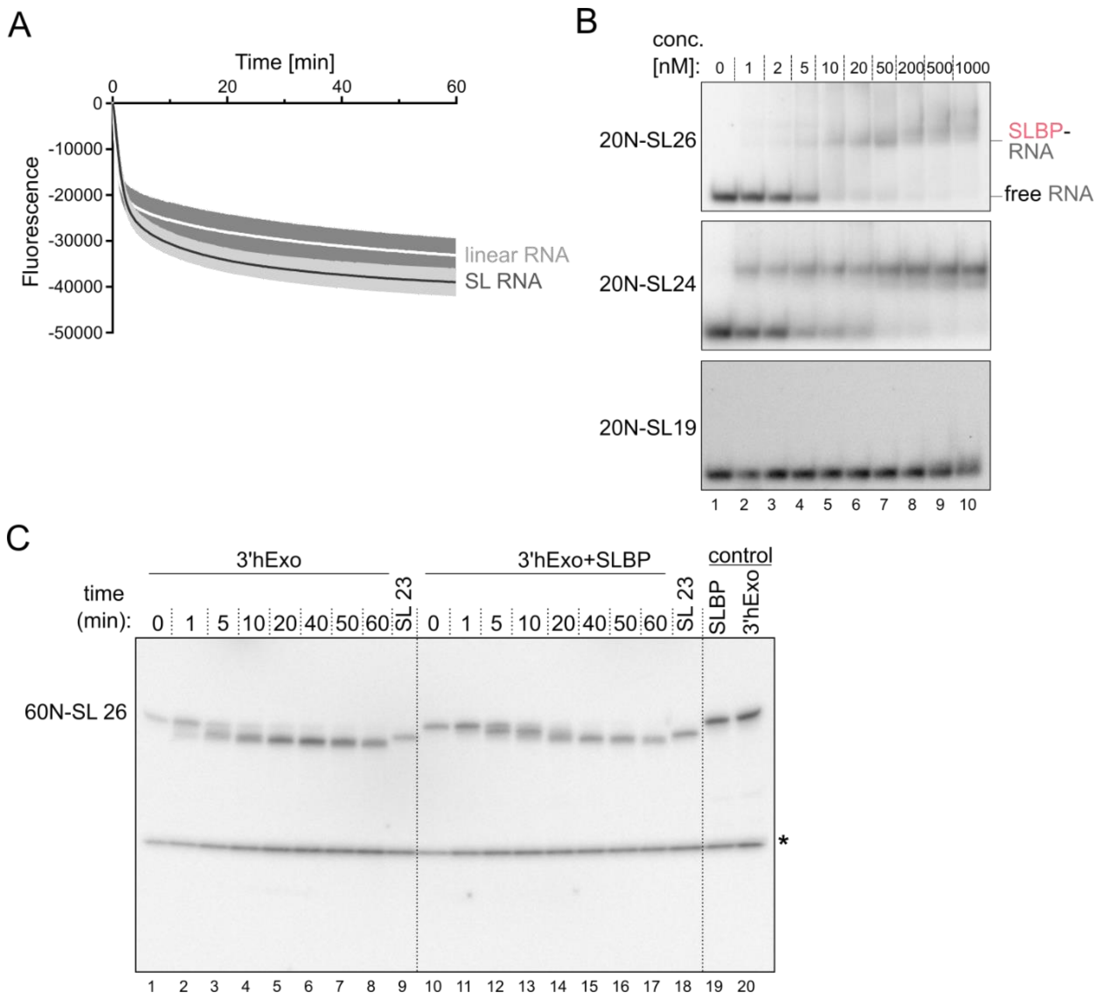


**Supplementary figure 3** (related to Figure 4)

**(A)** Time-dependent measurements of UPF1-Hel unwinding activity on the linear (light grey) and SL RNA (black) substrates. The decrease in fluorescence is relative to the "no protein" sample.

**(B)** Electrophoretic mobility shift assay (EMSA) of different histone SL RNA intermediates with increasing concentrations of SLBPfl. The SL RNAs were labelled with $^{32}$P. Protein-RNA complexes were resolved on native-PAGE gels and visualized by phosphorimaging. SLBP binds SL26 and SL24 with very high affinity but shows no binding to SL19, corroborating our fluorescence anisotropy measurements (Figure 3F). Generation of the 20N-SL RNA substrates by in vitro transcription is outlined in Supplementary figure 4B.

**(C)** Time-dependent analysis of degradation of 60N-SL26 RNA by 3'hExo in the absence (lanes 1-8) and presence (lanes 10-17) of SLBP. The degradation reaction was initiated by addition of magnesium ions. Samples were removed from the reaction at the indicated time points and the reaction was stopped by addition of RNA loading buffer. The samples were analysed by urea-PAGE, followed by phosphorimaging. Degradation of SL26 stops at the base of the stem, after removal of 5 nucleotides from the 3'-end. Addition of SLBP slows down degradation of SL26, acting as a roadblock for 3’hExo (compare lanes 3 and 12). * denotes a labelled DNA loading control used in this experiment.

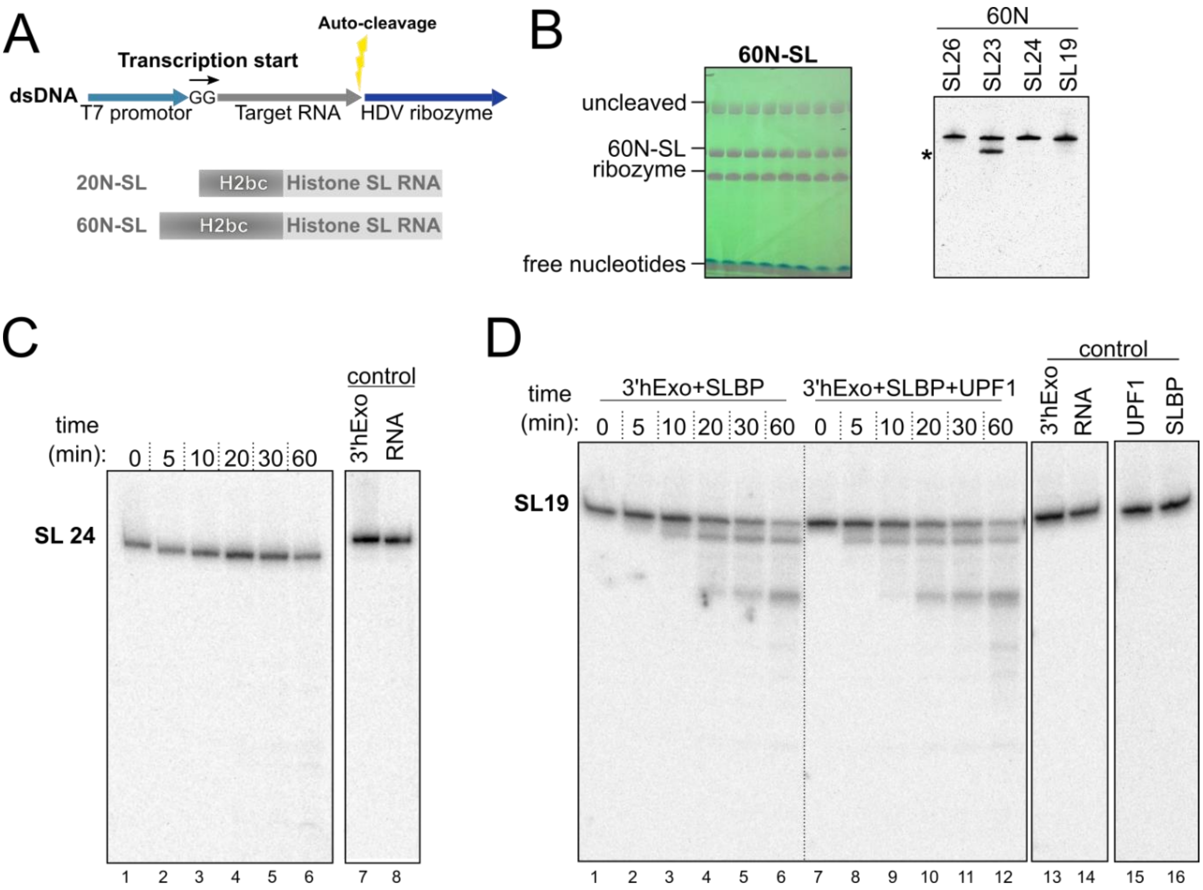


**Supplementary figure 4** (related to Figure 5)

**(A)** Schematic representation of the IVT template used to generate precise SL RNA species (top) and RNA substrates used for EMSA (20N-SL, middle) or *in vitro* degradation assays (60N-SL, bottom). Fusion of the HDV ribozyme to the DNA template sequence generates a precursor RNA that is spontaneously cleaved by the ribozyme to release SL RNA with precise 3'-ends. The sequence upstream of the conserved mammalian histone SL RNA is derived from the histone H2bc mRNA.

**(B)** Urea-PAGE analysis of IVT reactions of the precursor 60N-SL26-HDV RNA that is autocleaved to release the mature 60N-SL26 RNA (left panel). Bands on the gel were visualized by UV-shadowing. Cis-cleavage by the ribozyme generates precise 3'-ends at the end of the SL for each RNA, which cannot be ensured by transcription termination of RNA polymerase. The top, middle and lower bands correspond to the uncleaved product, the mature 60N-SL26 RNA and the ribozyme, respectively. The right panel shows a urea-PAGE analysis of all RNA substrates generated using this method. SL23 shows a doublet, arising from an artefact of IVT. For biochemical studies, a single homogenous species, corresponding to the top band, is obtained by further gel purification

**(C)** Time-dependent analysis of degradation of 60N-SL24 RNA by 3'hExo. The reaction was performed and analysed as described for Figure 4A. Degradation of SL23 stops at the base of the stem, after removal of 2 nucleotides from the 3'-end.

**(D)** Time-dependent analysis of degradation of 60N-SL19 RNA by 3'hExo in the presence of SLBPfl (lanes 1-6) and additionally in the presence of UPF1 (lanes 7-12). Lanes 13-16 represent negative controls to check for presence of nuclease contaminants in the proteins used. The negative control for 3'hExo lacked magnesium ions that are necessary for 3'-5' ribonucleolytic activity of 3'hExo. Addition of UPF1 enhances the efficiency of degradation by 3'hExo, as seen for Figure 4A.

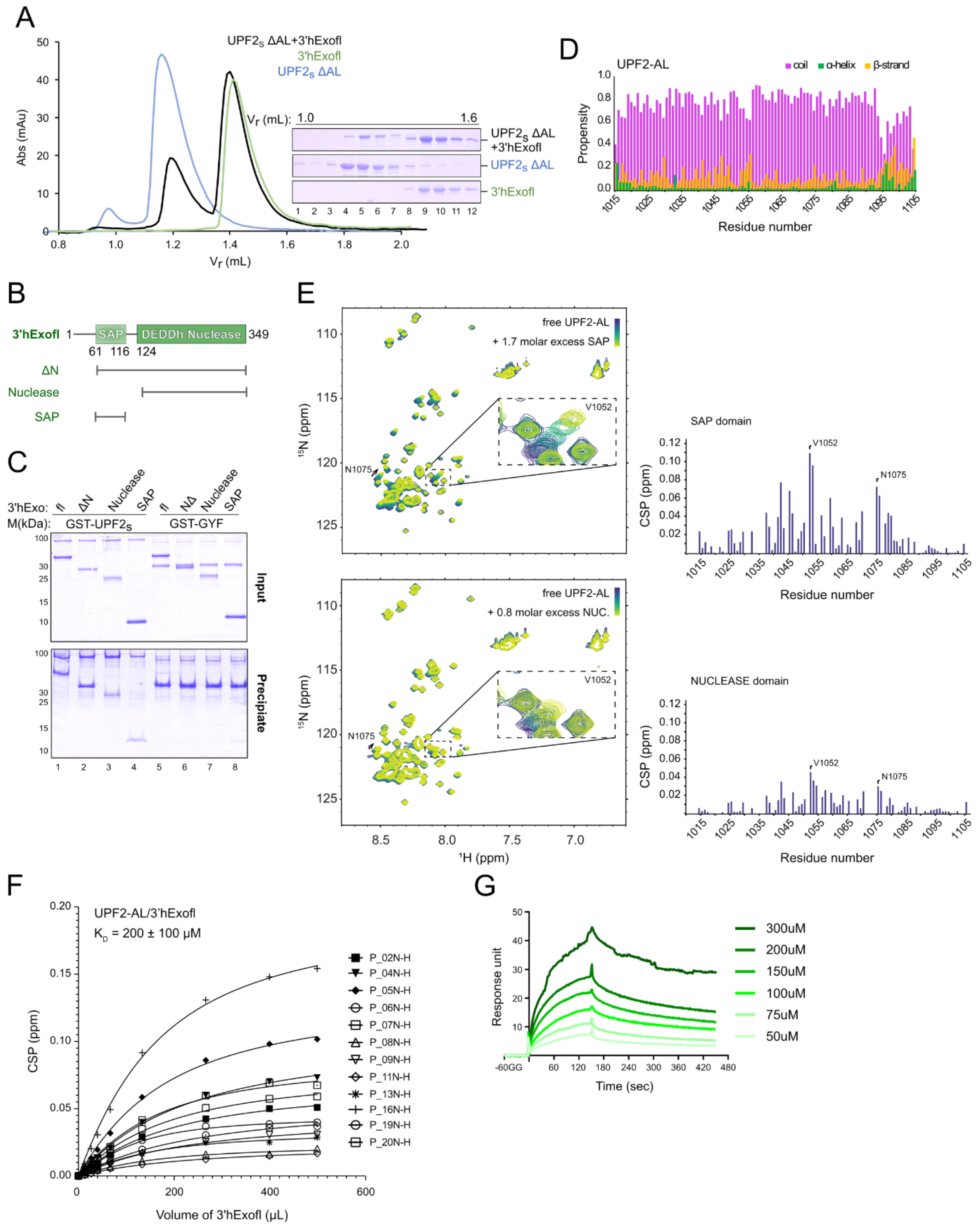


**Supplementary figure 5** (related to Figure 6)

**(A)** Analytical SEC of 3'hExofl and a UPF2 protein lacking the acidic linker (UPF2ΔAL). No complex is obtained in this case, confirming that the acidic linker of UPF2 is necessary for binding to 3'hExo.

**(B)** Schematic representation of the domain organization of human 3'hExo. Protein variants used in this study are shown below. Residue numbers indicate domain boundaries.

**(C)** GST-pulldown assay of GST-UPF2s (and GST-GYF as negative control) with 3'hExo variants. Both SAP and nuclease subdomains of 3'hExo bind UPF2.

**(D)** Secondary structure prediction of UPF2-AL based on NMR chemical shifts.

**(E)** $^{1}H$-$^{15}N$-HSQC NMR titration experiments of $^{15}N$-labeled UPF2s with increasing concentrations of 3'hExo-SAP and Nuclease subdomains. The spectrum of free UPF2-AL is in blue while those recorded in presence of increasing concentrations of 3'hExo proteins are in progressively lighter shades of green. The insets show a zoomed-in view of residue V1052 of UPF2-AL that show the largest chemical shift perturbations (CSP) upon addition of 3'hExo-SAP and Nuclease proteins, as observed with 3′hExofl (Figure 6F). The histograms of the CSPs of UPF2-AL upon titration of 3'hExo-SAP and Nuclease are plotted against the UPF2-AL protein sequence and shown below each spectrum. The patterns of CSPs obtained upon titration of the SAP and Nuclease subdomains are very similar, suggesting that they bind the same stretch of residues of UPF2.

**(F)** Plot of CSP vs. volume of 3'hExofl added to derive the dissociation constant ($K_D$) of the UPF2-AL:3'hExo interaction. To ensure that the concentration of UPF2-AL protein could be accurately determined and its purity assessed by SDS-PAGE analysis, a modified version of the UPF2 acidic linker was designed spanning residues 1024-1085 and containing three lysine residues and one tryptophan at the N- and C-termini, respectively.

**(G)** Binding affinity of 3'hExofl for UPF2-AL1 as measured by surface plasmon resonance. 3'hExofl was immobilized on a CM5 chip and UPF2-AL1 was injected at concentrations ranging from 50-300 μM. Binding constants were derived from 3 independent experiments by fitting the data to a 1:1 binding model.

### Supplementary Table 1

| **DNA oligonucleotide templates for IVT** | | |
|---|---|---|
| **Oligo Name** | **Description** | **Sequence (5'→3')** |
| oAA28 | 60N-SL+HDV oligo fwd | CTAATACGACTCACTATAGGCCGTCACCAAGTAC |
| oAA54 | 60N-SL26+HDV oligo rev | GTCCCATTCGCCATCGCGAACGATGTTGCCCACCGGCCG<br>CCAGCGAGGAGGCTGGGACCATGGCCGGCTGGGTGGCTC<br>TGAAAAGAGCCTTTGGGGTTAG |
| oAA55 | 60N-SL19+HDV oligo rev | GTCCCATTCGCCATCGCGAACGATGTTGCCCACCGGCCG<br>CCAGCGAGGAGGCTGGGACCATGGCCGGCCTCTGAAAAG<br>AGCCTTTGGGGTTAG |
| oAA26 | 60N-SL23+HDV oligo fwd | CTAATACGACTCACTATAGG<br>CCGTCACCAAGTACACCAGCTCCAAGTAAACATTCCAAG<br>TAAGCGTCTTAACACCTAACCccaaaggctctttt |
| oAA27 | 60N-SL23+HDV oligo rev | CTAACCccaaaggctcttttcagag ccac<br>GCCGGCCATGGTCCCAGCCTCCTCGCTGGCGGCCGGTGG<br>GCAACATCGTTCGCGATGGCGAATGGGAC |
| oAA118 | 60N-SL24+HDV oligo rev | GTCCCATTCGCCATCGCGAACGATGTTGCCCACCGG<br>CCGCCAGCGAGGAGGCTGGGACCATGGCCGGCAGTG<br>GCTCTGAAAAGAGCCTTTGGGGTTAGGT |
| oAA79 | SL RNA unwinding assay oligo fwd | CTAATACGACTCACTATAGGGACACAAAACAAAAGACAA<br>AAACACAAAACAAAAGACAAAAACACAAAACAAAAG |
| oAA80 | SL RNA unwinding assay oligo rev | AGCTAGTTGTACGCACACGGTGGCTCTGAAAAGAGCCTT<br>TGGCTTTTTGTCTTTTGTTTTGTGTTTTTGTCTTTT |
| oAA81 | Linear RNA unwinding assay oligo rev | AGCTAGTTGTACGCACACGGTAATTTGGCTTTTTGTCTT<br>TTGTTTTGTGTTTTTGT |
| **RNA substrates generated by IVT** | | |
| **Name** | **Description** | **Sequence (5'→3')** |
| 60N-SL26 | upstream H2bc sequence(60 nt) + histone SL 26mer (full-length) | GGCCGUCACCAAGUACACCAGCUCCAAGUAAACAUUCCA<br>AGUAAGCGUCUUAACACCUAACCCCAAAGGCUCUUUUCA<br>GAGCCACCCA |
| 60N-SL23 | upstream H2bc sequence(60 nt) + histone SL 23mer (FL-3 nt) | GGCCGUCACCAAGUACACCAGCUCCAAGUAAACAUUCCA<br>AGUAAGCGUCUUAACACCUAACCCCAAAGGCUCUUUUCA<br>GAGCCACU |
| 60N-SL24 | upstream H2bc sequence(60 nt) + histone SL 24mer (FL-3 nt+1 U) | GGCCGUCACCAAGUACACCAGCUCCAAGUAAACAUUCCA<br>AGUAAGCGUCUUAACACCUAACCCCAAAGGCUCUUUUCA<br>GAGCCAC |
| 60N-SL19 | upstream H2bc sequence(60 nt) + SL 19mer (FL-7nt) | GGCCGUCACCAAGUACACCAGCUCCAAGUAAACAUUCCA<br>AGUAAGCGUCUUAACACCUAACCCCAAAGGCUCUUUUCA<br>GAG |
| SL UWA | for unwinding assay, SL 24mer (FL-2 nt), downstream hybridization site (18 nt), upstream UPF1 translocation site (63 nt) | GGGACACAAAACAAAAGACAAAAACACAAAACAAAAGAC<br>AAAAACACAAAACAAAAGACAAAAAGCCAAAGGCUCUUU<br>UCAGAGCCACCGUGUGCGUACAACUAGCU |

| | | |
|---|---|---|
| Linear UWA | for unwinding assay, ss ctrl substrate, downstream hybridization site (18 nt), upstream UPF1 translocation site (63 nt) | GGGACACAAAACAAAAGACAAAAACACAAAACAAAAGACAAAAACACAAAACAAAAGACAAAAAGCCAAAUUACCGUGUGCGUACAACUAGCU |

**Synthetic RNAs**

| Name | Sequence (5'→3') |
|---|---|
| 15U-26SL | UUUUUUUUUUUUUUUCCAAAGGCUCUUUUCAGAGCCACCCA |
| 5‘ 6-FAM-SL26 | 6-FAM/CCAAAGGCUCUUUUCAGAGCCACCCA |
| 5‘ 6-FAM-SL24 | 6-FAM/CCAAAGGCUCUUUUCAGAGCCACU |
| 5‘ 6-FAM-SL19 | 6-FAM/CCAAAGGCUCUUUUCAGAG |
| 5‘ Alexa488-labeled DNA probe | Alexa488/AGCTAGTTGTACGCACAC |
| 3‘ Black Hole Quencher 1- conjugated DNA trap | GTGTGCGTACAACTAGCT/3BHQ_1- 3’ |

**Primer and sgRNA Sequences**

| Name | Sequence (5'→3') |
|---|---|
| SLBP-gF1 | CCG GGG ACG CGG TCG GCT GGG CAC |
| SLBP-gR1 | AAA CGT GCC CAG CCG ACC GCG TCC |
| SLBP-gF2 | CCG GCG AGG GAG CGC GTG CCC CGT |
| SLBP-gR2 | AAA CAC GGG GCA CGC GCT CCC TCG |
| SLBP-gF3 | CCG GTC GCG GTG CCG GGA TCG GTC |
| SLBP-gR3 | AAA CGA CCG ATC CCG GCA CCG CGA |
| SLBP-gF4 | CCG GGG AGC CCG CGG CCT CGT CAA |
| SLBP-gR4 | AAA CTT GAC GAG GCC GCG GGC TCC |
| SLBP-tF | GTTGTAAGGCGGTCCCGAA |
| SLBP-tR | GTTCAGCTGACTCCCATCCT |
| SLBP-RT-F | ATCAGAGCCGCTGCGACGGTGACGCCA |
| SLBP-RT-R | TCAGAACTTCCTGATGATGA |

**Supplementary Methods**

*RNA radiolabelling and Electrophoretic mobility shift assays (EMSA)*

Prior to radiolabelling, RNA samples were dephosphorylated at their 5' end using the QuickCIP enzyme (NEB). 10 pmol of RNA was radiolabelled at the 5′-end with [γ-$^{32}$P]-ATP using T4 Polynucleotide Kinase (T4PNK) (ThermoFisher). The labelled RNA was separated from the excess radiolabeled ATP by gel purification from a 10 % denaturing PAGE. The gel fragment corresponding to the radiolabeled substrate was excised and gel pieces were crushed and soaked overnight in G50 buffer (20 mM Tris-HCl, 300 mM sodium acetate, 2 mM EDTA, 0.25 % SDS). After phenol/chloroform extraction and ethanol precipitation, RNA was dissolved in nuclease-free water. The $^{32}$P-labeled substrates were stored at −20 °C until further use.

For EMSAs, 0.2 pmol of radiolabelled RNA was mixed with varying amounts of full-length SLBP in EMSA binding buffer (20 mM HEPES, 100 mM NaCl, 0.1 % NP-40, 0.5 % Glycerol, 1 mM DTT, 0.5 mM EDTA) supplemented with 100 ng tRNA in a total volume of 10 µL. The samples were incubated on ice for 1 h and then resolved on a 5 % native PAGE. The labelled RNA was visualized by phosphorimaging after exposure of the gel to an image plate (Cytiva).

*Surface Plasmon Resonance*

Surface plasmon resonance experiments were performed on a Biacore Upgrade instrument (Biacore™ X100) at 25 °C. Research grade carboxymethyldextran (CM5) chips were purchased from Cytiva. *N*-hydroxysuccinimide (NHS), *N*-ethyl-*N*′-(3-dimethylaminopropyl) carbodiimide hydrochloride (EDC) and ethanolamine were also procured from Cytiva. HBS-EP (10 mM HEPES pH 7.5, 125 mM NaCl, 3 mM EDTA, 0.005% Biacore Surfactant P20) and 10 mM sodium acetate buffer were prepared in lab. The CM5 sensor Chip was soaked in HBS-EP buffer overnight at room temperature before use. Then the chip was dried and loaded to the Biacore chamber. 3'hExoI was diluted to 1.2 µg/mL with 10 mM sodium acetate pH 5.0 buffer and injected in pulses on an EDC/NHS activated CM5 Biacore chip until 1900 response units (RU) were immobilized on the CM5 Biacore chip. Remaining activated carboxyl groups were quenched with Ethanolamine. The UPF2-AL protein, ranging from 300 μM to 50 μM, were then applied onto the chip and injected for a contact time of 150 s. 5 M NaCl was used as a regeneration solution. All experiments were repeated three times. The $k_{on}$, $k_{off}$ and $K_D$, were determined by analyzing the kinetic data by a global fit to a 1:1 binding model with the BIA evaluation software (Biacore, Cytiva).
